# Unconventional stabilization of the human T-cell leukemia virus type 1 immature Gag lattice

**DOI:** 10.1101/2023.07.24.548988

**Authors:** Martin Obr, Mathias Percipalle, Darya Chernikova, Huixin Yang, Andreas Thader, Gergely Pinke, Dario Porley, Louis M Mansky, Robert A Dick, Florian KM Schur

## Abstract

Human T-cell leukemia virus type 1 (HTLV-1) has an atypical immature particle morphology compared to other retroviruses. This indicates that these particles are formed in a way that is unique. Here we report the results of cryo-electron tomography (cryo-ET) studies of HTLV-1 virus-like particles (VLPs) assembled *in vitro*, as well as derived from cells. This work shows that HTLV-1 employs an unconventional mechanism of Gag-Gag interactions to form the immature viral lattice. Analysis of high-resolution structural information from immature CA tubular arrays reveals that the primary stabilizing component in HTLV-1 is CA-NTD. Mutagenesis and biophysical analysis support this observation. This distinguishes HTLV-1 from other retroviruses, in which the stabilization is provided primarily by the CA-CTD. These results are the first to provide structural details of the quaternary arrangement of Gag for an immature deltaretrovirus, and this helps explain why HTLV-1 particles are morphologically distinct.

## Introduction

The *Retroviridae* family includes two important human pathogens infecting T cells, human immunodeficiency virus type 1 (HIV-1) and human T-cell leukemia virus type 1 (HTLV-1). The global prevalence suggest that the number of people living with HTLV-1 (a member of the *Deltaretrovirus* genus) ranges from 5-10 million, which is likely an underestimate (*1*). While most HTLV-1 infections remain asymptomatic, approximately 5% lead to aggressive diseases such as HTLV-1-associated myelopathy/tropical spastic paraparesis (HAM-TSP) or adult T-cell leukemia/lymphoma (ATLL). ATLL is an aggressive form of cancer with a median survival rate of less than one year (*2*, *3*).

The retroviral structural protein Gag forms the immature protein shell of nascent virus particles (*4*). All retroviral Gag proteins contain three canonical domains (**Figure S1A**): matrix (MA), capsid (CA) – consisting of independently folded N-terminal CA (CA-NTD) and C-terminal CA (CA-CTD) domains – and nucleocapsid (NC). During immature virus particle formation, these domains function in membrane binding (MA), viral lattice self-assembly (CA), and viral genomic RNA (vgRNA) packaging (NC), respectively. Gag oligomerization is primarily driven by interactions between CA domains, and these interactions determine virus particle morphology and size (*5*). The immature Gag lattice has local six-fold symmetry and is incomplete; e.g. in HIV-1 it covers ∼60% of the available membrane surface area inside a virion. Upon maturation, the viral-encoded protease cleaves Gag at defined positions, leading to a cascade of structural changes that rearrange the virion interior(*6*). In a mature virion MA remains associated with the viral membrane while the CA protein forms a capsid core consisting of CA hexamers and pentamers. The capsid core contains the condensed NC-vgRNA complex, reverse transcriptase, and integrase. *In vitro* expression of Gag in mammalian cells, and bacterial expression and purification of certain truncated variants thereof, is sufficient for assembly of virus-like particles (VLPs) *in vitro* (*7*) which have authentic immature Gag or mature CA architectures (*8–11*).

Despite substantial sequence variation among retrovirus species, CA shows a strongly conserved tertiary fold, with six to seven **α**-helices in the CA-NTD and four **α**-helices in the CA-CTD (*12–18*). The latter harbors the highly conserved major homology region (MHR), which has been implicated in preserving the CA-CTD protein fold and in establishing relevant interactions in the assembly of retroviral lattices (*9*, *19*). HIV-1 (*Lentivirus*) and Rous sarcoma virus (RSV, *Alpharetrovirus)* have a Gag cleavage product between CA and NC named spacer peptide (SP), and Mason-Pfizer monkey virus (M-PMV, *Betaretrovirus)* has a similar segment in CA named the spacer peptide-like region (*20*, *21*). The SP region of Gag is important for immature assembly and in the regulation of maturation(*4*). Similarly, Gag and CA lattice formation in immature and mature virus particles, respectively, requires CA-CTD dimerization, which is established via hydrophobic residues in helices 8 or 9 of the CA-CTD (*22*).

The CA-CTD has been proposed to have a dominant role for immature assembly of retroviruses. Variants of HIV-1 Gag containing only the C-terminal half of CA and downstream regions can form VLPs when expressed in cells (*23*, *24*). Cryo-electron tomography (cryo-ET) and subtomogram averaging of purified immature virus particles or *in vitro*-assembled immature-like VLPs of lentiviruses, and alpha-, beta-, and gamma-retroviruses demonstrate that CA-CTD dimerization is a key factor in the conserved structural arrangement of the CA-CTD hexamer of immature Gag lattices (*8*, *18*, *25*, *26*). These studies also show that the immature CA-NTD arrangement differs significantly between retrovirus genera. In summary, these structural studies are consistent with the hypothesis that the CA-CTD is responsible for forming the essential stabilizing immature assembly interfaces in retroviruses, while the CA-NTD may play a primary role in determining immature lattice curvature and particle size and representing a major binding site for host cell factors (*27*).

It is not certain if the function of the CA-NTD in assembly is conserved for retroviruses, and if there are other CA-NTD arrangements which have not been reported. Interestingly, previous mutagenesis experiments targeting residues in the CA-CTD of HTLV-1 Gag did not affect VLP budding (*28*), suggesting that unlike other retroviruses, the CA-CTD is not the key determinant of immature assembly. Morphological studies of HTLV-1 particles using cryo-electron microscopy (cryo-EM) also showed peculiar differences in Gag lattice shape, such as areas of flat lattices and varying membrane-CA distance (*29–31*). Taken together these studies suggest that HTLV-1 may employ different Gag assembly mechanisms than other retroviruses.

A comparison of the structural details of the immature HTLV-1 viral lattice to other retroviruses would provide insight on the structural conservation and diversity of retroviral Gag assembly mechanisms. However, there is currently a lack in structural information for HTLV-1. Given that HTLV-1 is a human pathogen of significance, and the ongoing development of inhibitors targeting HIV-1 CA (capsid assembly inhibitors) (*32*), it is critical to generate this structural information.

Here we report cryo-ET and subtomogram averaging results of full-length and truncated HTLV-1 Gag assemblies (**Figure S1A-B**). We describe the key immature HTLV-1 assembly determinants. Furthermore, our supporting mutagenesis experiments and biophysical characterization delineate the contributions of the CA-NTD and CA-CTD to assembly. Our results show that HTLV-1 employs a novel CA-NTD arrangement to form the immature Gag lattice and that, in contrast to other retroviral structures, the CA-NTD is the key determinant of immature HTLV-1 assembly.

## Results

### HTLV-1 Gag VLPs display unusual variability

To gain insight into the immature architecture of HTLV-1 we employed cryo-ET to visualize mammalian cell-derived Gag-based VLPs (**Table S1, Figure 1A, Figure S1C, Movie S1**). As previously reported (*30*, *31*), the shape and size of the VLPs varied substantially. The majority of VLPs were spherical, however, the local curvature ranged from flat patches to sharp kinks (**Figure 1A, Movie S1**). As with other retroviruses, the individual domains of Gag formed distinct layers of density starting at the inner leaflet of the viral membrane (corresponding to MA) and extending towards the center of the VLP by ∼25 nm (**Figure 1B-C**). The CA-layer exhibited a regular organization and so we employed subtomogram averaging (**Figure S2A**) in order to obtain higher resolution information for this layer. Alignments focused on the CA layer resulted in a blurred density for the membrane, which is consistent with a measured variation in the CA-to-membrane distance (mean distance of 17.7 nm, standard deviation ±0.8nm) (**Figure S2B-C**). The distance of flat Gag lattice regions to the membrane was previously reported to be greater than the distance at curved regions (*30*).

**Figure 1:**
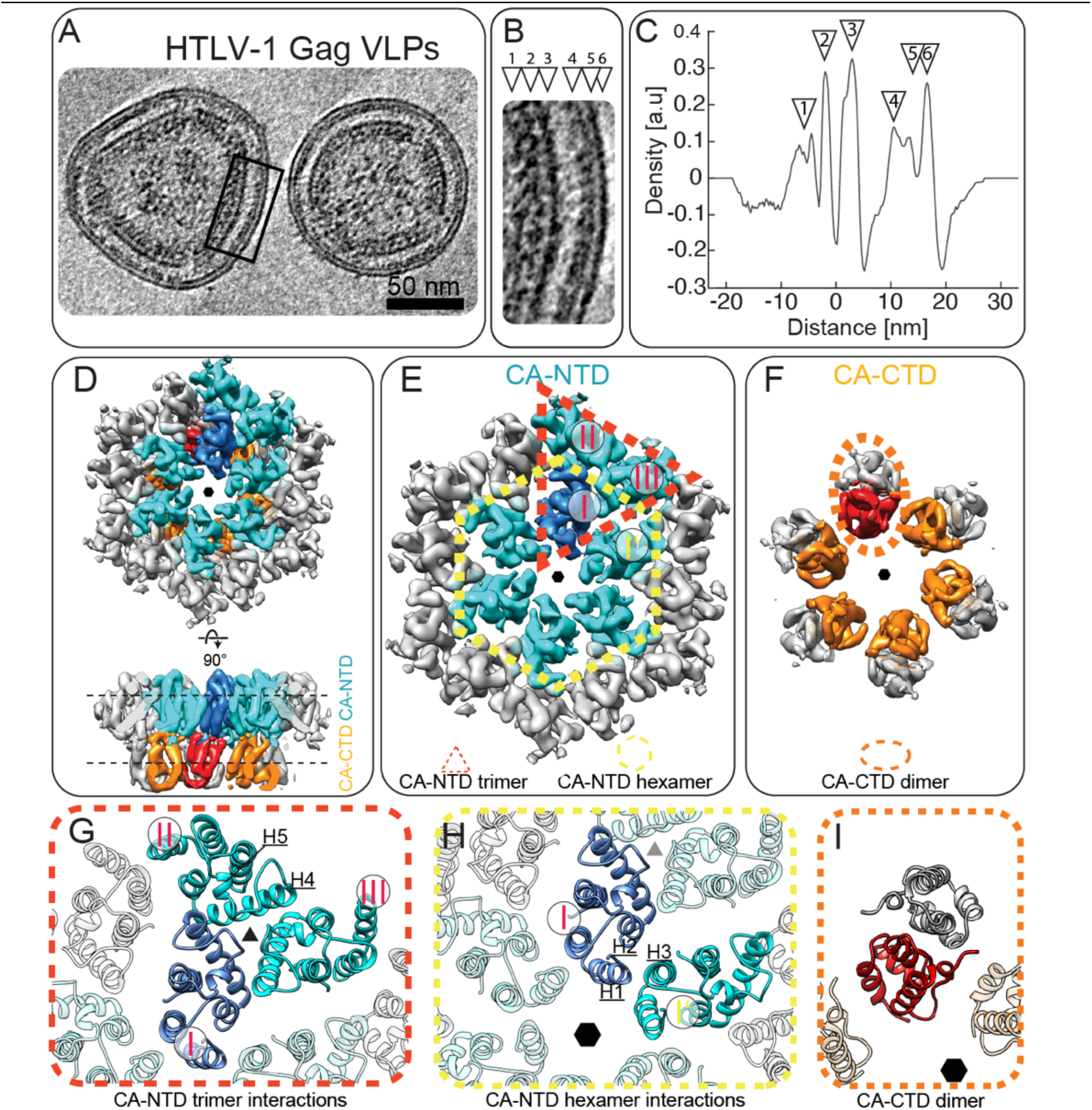
Cryo-ET of immature HTLV-1 Gag VLPs. **A)** Computational slice (8.8 nm thickness) through a cryo-electron tomogram containing HTLV-1 Gag-based VLPs. Protein density is black. Scale bar is 50 nm. **B)** Enlarged view of the Gag lattice within VLPs, as annotated by a black rectangle in panel (A). The arrowheads designate the different layers of the radially aligned Gag lattice underneath the viral membrane. NC-RNP (1), CA-CTD (2), CA-NTD (3), MA (4), inner leaflet (5), outer leaflet (6). **C)** 1D radial density plot of the Gag lattice in immature HTLV-1 Gag-based VLPs, measuring the distance of the individual Gag domains and the viral membrane from the linker between the CA-NTD and CA-CTD. The annotation with arrowheads is as in panel (B). a.u. arbitrary units. **D-F)** Isosurface representations of the subtomogram average of the CA hexamer from HTLV-1 Gag-based VLPs. The CA-NTDs of the central hexamer are colored cyan, with one monomer highlighted in blue. Two additional CA-NTD monomers from adjacent hexamers are also colored in cyan, to highlight the trimeric inter-hexamer interface. The CA-CTDs of the central hexamer are colored orange/red. The hexameric arrangement is indicated by a small hexagon. **D)** CA lattice as seen from the outside of the VLP and rotated by 90 degrees to show a side view. **E)** Top view of the CA-NTD, with the trimeric inter-hexamer interface and the intra-hexameric interfaces indicated with a dashed red triangle and dashed yellow hexamer, respectively. CA monomers are annotated as described in the main text. **F)** Top view of the CA-CTD hexamer. A dashed orange ellipsoid highlights one CA-CTD dimer linking adjacent hexamers. **G-H)** Molecular models of the CA-NTD and CA-CTD rigid body fitted into the EM-density of the immature CA lattice. Coloring as in D-F. **G)** Trimeric CA-NTD interactions linking hexamers. **H)** Interactions around the hexamer, involving helices 1,2 and 3 from adjacent CA-NTDs. **I)** Model of the CA-CTD dimer.

Retroviral Gag lattices are spherical and the CA-layer follows local C6 symmetry (*8*, *18*, *33*). However, to account for the observed heterogeneity of HTLV-1 Gag VLP shapes, we applied local C2 symmetry during subtomogram averaging. The resulting maps had a CA-CTD layer that was resolved to a lower resolution than the CA-NTD layer, which we predict is due to heterogeneity caused by flexibility of the CA-CTD. 3D classification focused on the CA-CTD yielded a class in which all helices were resolved. In contrast, the CA-NTD did not show increased heterogeneity. Therefore, we used both domains as independent species in a multiparticle refinement using the software M (*34*). This approach yielded 5.9 and 6.2Å resolution maps of the immature CA-NTD and CA-CTD layers, respectively (**Figure 1D-F**, **Figure S2D**). This allowed for the generation of a model of the immature HTLV-1 CA lattice by rigid body fitting models of HTLV-1 CA-NTD and CA-CTD into the EM-density map. Densities for MA and NC above and below CA, respectively, are present in the 1D density plot (**Figure 1C**), but MA and NC are not resolved in the subtomogram averages, suggesting that neither domain follows the same organization as the CA lattice.

### Interactions across two CA-NTD interfaces stabilize the immature lattice

The CA-NTD establishes lateral interactions that form two basic building blocks shaping the lattice – a CA-NTD trimer and a CA-NTD hexamer (**Figure 1E, 1G, 1H, Movie S2**). For both the trimer and hexamer, each CA-NTD monomer contacts two adjacent monomers around the local pseudo-symmetry axes. Specifically, the trimer is formed by interactions spanning residues in helices 4 and 5 (**Figure 1E, G**). Similarly, the hexamerization interface involves helices 1, 2 and 3 of two neighboring monomers (annotated I and I’ in **Figure 1E and 1H**). Comparison to other immature retroviral CA arrangements shows that HTLV-1 adopts an entirely different CA-NTD arrangement within its immature lattice (**Figure S3**). Unlike other retroviruses (*8*, *18*, *25*), there is no dimerization interface in the HTLV-1 CA-NTD. Instead, there is a large gap at the local C2 symmetry axis, lined with hydrophilic residues of helices 2, 4, and 6.

### CA-CTD forms isolated dimers, but not a continuous lattice

Within the CA-CTD layer, the only intermolecular contact is a dimerization interface formed by helix 8 (**Figure 1F, 1I**, **Figure S4A, Movie S2**). This is similar to other immature retroviral Gag assemblies, where contacts between hexamers are established via a dimeric interface involving two neighboring CA-CTD monomers (**Figure S4A**). Unlike other retroviruses, where the CA-CTD forms inter-and intra-hexameric contacts, the CA-CTD of HTLV-1 only forms CA-CTD inter-hexameric contacts. The positioning of the individual HTLV-1 CA-CTD dimers places them out of reach of the adjacent dimers around the hexameric ring (**Figure S4B**), not enabling them to contribute to intra-hexamerization interfaces forming a lattice.

HTLV-1 lacks a six-helix bundle (6HB) which is reported to be critical for immature CA hexamer formation for other retroviruses. For HIV-1 the 6HB is formed by residues in CA and SP1 (*9*, *35*). Similarly, in RSV, M-PMV, and murine leukemia virus (MLV) the immature lattices are stabilized by interactions downstream of the CA-layer via a 6HB, which are clearly observed from cryo-ET and subtomogram averaging of these retroviruses (*8*, *18*, *25*) (**Figure S4B**). HTLV-1 Gag contains only the canonical Gag domains, MA, CA, and NC, and is not reported to have a spacer peptide between CA and NC. This is consistent with our observation that there are no ordered densities C-terminal to CA, which would be indicative of the presence of a 6HB or another stabilization-providing region. Hence, the absence of an assembly element like a 6HB may explain why HTLV-1 has a less well-organized CA-CTD layer.

The HTLV-1 CA-CTD effectively acts to help stabilize the CA-NTD inter-hexameric interactions. The positions of individual dimers are constrained only by the flexible linker between CA-NTD and CA-CTD. The role of CA-CTD seems to be in further crosslinking the NTD layer, while not constraining the shape of the lattice to a single curvature. Instead, the variable distance between the CA-CTD dimers allows for heterogeneity in particle size and shape.

### A high-resolution model of CA-NTD reveals the architecture of the immature HTLV-1 lattice

To analyze the structure of immature HTLV-1 lattice interfaces in more detail, we engineered an HTLV-1 Gag truncation construct for bacterial expression, purification, *in vitro* assembly and analysis by cryo-ET. Our construct lacked the N-terminal 125 residues of Gag (MA126CANC) (**Figure S1A, S5A**), similar to other Gag truncation variants that have been previously used to study immature HIV-1, equine infectious anemia virus (EIAV) and M-PMV (*9*, *20*, *26*).

The purified HTLV-1 MA126CANC assembled into hollow tubes (**Figure 2A, Movie S3**), similar to immature-like tubes observed for EIAV, M-PMV and HIV-1 (*20*, *26*, *36*). Cryo-ET of the HTLV-1 tubular assemblies revealed a density profile of the CANC layer matching the profile observed for HTLV-1 Gag-based VLPs (compare **Figure 1C and 2B**). The MA126CANC tubes exhibited different helical parameters. Tubes were sorted into nine groups and each group was processed separately by subtomogram averaging (**Figure S6**). The two groups with the greatest number of tubes had similar geometry, and so they were pooled and processed together. The resulting final reconstruction of the CA-NTD layer had an overall estimated resolution of 3.4 Å (**Figure S7A**). The helical pitch and densities for large side chains were clearly visible at this resolution, which enabled us to build a refined model of the immature CA-NTD lattice interactions (**Figure S7B**). The *in vitro*-assembled tubes (**Figure 2C**) had the same arrangement of the CA-NTD layer as the full-length HTLV-1 Gag VLPs (RMSDC**α**=1.2 Å), underscoring the biological relevance of our *in vitro* system. Hence, our refined model allowed identification of the side-chains likely to participate in the CA-NTD interfaces that stabilize the immature lattice.

**Figure 2:**
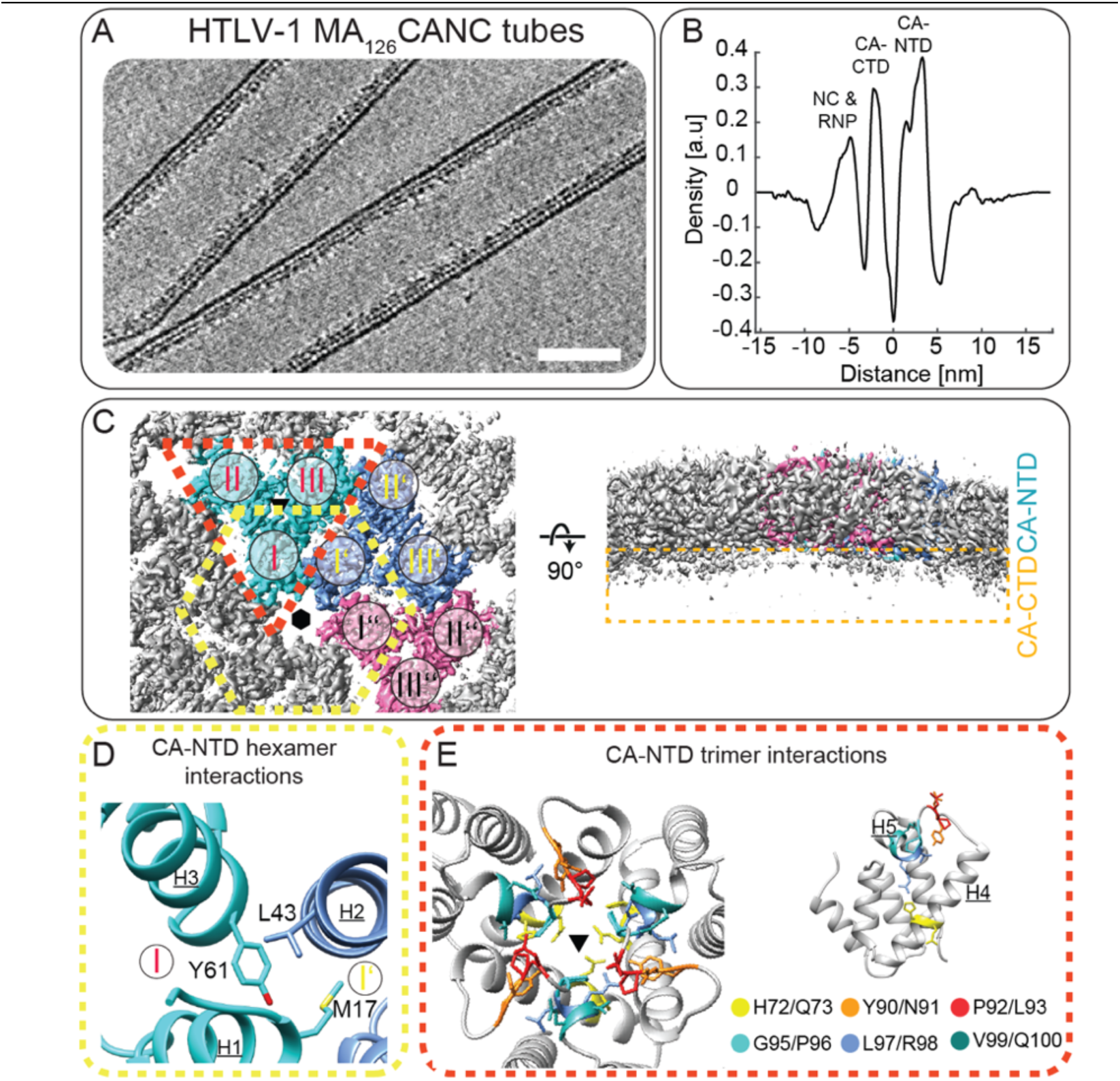
Structural model of the immature HTLV-1 CA-NTD interactions. **A)** Computational slice (8.8 nm thickness) through a cryo-electron tomogram containing HTLV-1 MA126CANC tubes. Protein density is black. Scale bar is 50 nm. **B)** 1D radial density plot of the CANC lattice in MA126CANC tubes, measuring the distance of the individual CA domains and NC-RNP from the linker between the CA-NTD and CA-CTD. **C)** EM density map of the immature HTLV-1 CA-NTD hexamer at 3.4 Å resolution, seen from the outside of the tube (left) and a side view (right). The three symmetry-independent trimer positions are colored in cyan, blue and pink. The remaining three CA-NTD domains of the central hexamer are colored in dark grey. Note the missing density for the CA-CTD in the side view on the right. **D)** Zoom-in view into the CA-NTD intra-hexamer interface (annotated with a yellow dashed hexagon in (C)). Labeling as in Figure 1. Assembly-relevant residues and the corresponding helices are annotated. **E)** Zoom-in view into the trimeric CA-NTD interface (annotated with a red triangle in (C)). Residues within the trimeric interface, which have been analysed via mutagenesis experiments are shown and colored according to the indicated coloring scheme. For simplicity one CA-NTD domain with the highlighted and colored residues is shown in side view on the right, to allow an easier appreciation of the residue location within the CA-NTD. The C2-hexamer center of the isosurface view in (C) is annotated by a schematic hexamer and the trimeric interfaces in (C) and (E) are annotated by black triangles.

### Residues involved in stabilizing immature CA-NTD lattice interactions

Previous tissue culture-based mutagenesis studies of the CA-NTD revealed the critical role of this domain for particle morphogenesis, while CA-CTD mutations did not affect budding. Specifically, CA-NTD residues M17 and Y61 have been identified as essential for immature HTLV-1 Gag lattice assembly and particle formation (*37*). In our model these residues are located in helices 1 and 3, respectively, which together with helix 2 form the hexamerization interface (**Figure 2D**). To further characterize this interface, we mutated residue L43, which we predicted based on our model, interacts with M17 and Y61. The L43A mutation reduced particle release by more than 50%, and further morphological characterization of released L43A particles showed aberrant and scarce patches of Gag lattice underneath the viral membrane (**Figure S8A-B**).

Double-alanine swap experiments on a set of adjacent residues at the trimeric CA-NTD interface were performed to characterize this interface. Residue pairs H72/Q73, Y90/N91, P92/L93, G95/P96, L97/R98, V99/Q100 were mutated (**Figure 2E**) and screened for altered particle production efficiency. These mutagenesis experiments revealed a substantial role for several of these residues in immature Gag assembly and particle release (**Figure S8C**), further validating our structural data. Consistent with our results, the G95/P96 pair was shown in an insertion/substitution mutation approach to reduce particle formation (*28*).

### The organization of the CA-CTD layer allows it to adopt variable curvatures

The CA-CTD layer in the tubular arrays also was arranged in a similar way as in the VLPs produced from cells (**Figure 1F, Figure S7C**), forming a layer of disconnected dimers. However, in this case the qualitative difference between CA-NTD and CA-CTD was even more dramatic. The CA-CTD density in different individual tubes did not permit structural analysis other than coarse positioning of the CA-CTD dimer, as the CA-CTD layer was only visible at lower resolution (compare **Figure 2C and Figure S7C**). Nevertheless, this allowed us to assess large-scale changes in the CA layer between the different tube geometries. Unlike the CA-NTD layer, the CA-CTD layer was observed to have larger movement of its subdomains. While the distance between the adjacent symmetry-independent CA-NTD trimers was nearly constant (42.1-42.6 Å), the distance between symmetry independent CA-CTD dimers was skewed such that it was decreased in the direction of tube curvature (31.8 Å) and increased along the tube axis (35.6 Å) (**Figure S7D**). Whether there are different conformations of CA-CTD dimers, and to what extent these might contribute to the flexibility/fluidity of the layer, could not be discerned from our data.

## The CA-CTD shows increased stability but lower interaction potential compared to CA-NTD and full-length CA

Given the peculiar behavior of the CA-CTD within the immature lattice, we sought to further characterize full-length CA and its two domains separately using biophysical approaches. To this end, HTLV-1 CA, CA-NTD (residues 131-258) and CA-CTD (residues 259-344) (**Figure S5A**) were expressed in *E. coli* and purified. The CA, CA-NTD and CA-CTD proteins were subjected to differential scanning fluorimetry (nanoDSF), which monitors changes in intrinsic tryptophan fluorescence (ITF) upon protein unfolding as a function of temperature (*38*) (**Figure 3A-C**). We also conducted back-reflection and dynamic light scattering (DLS) experiments to measure turbidity and determine cumulant radius as a means of informing about aggregation and assembly properties (**Figure 3D-I**). The measurements were carried out in three different conditions: 1) with non-assembled protein in storage buffer, 2) with non-assembled proteins in assembly buffer (*i.e.,* inducing assembly just before measurement), and 3) with assembled protein in assembly buffer. We were able to obtain reproducible assemblies only for HTLV-1 CA (**Figure S5B)**, but not its domains.

**Figure 3:**
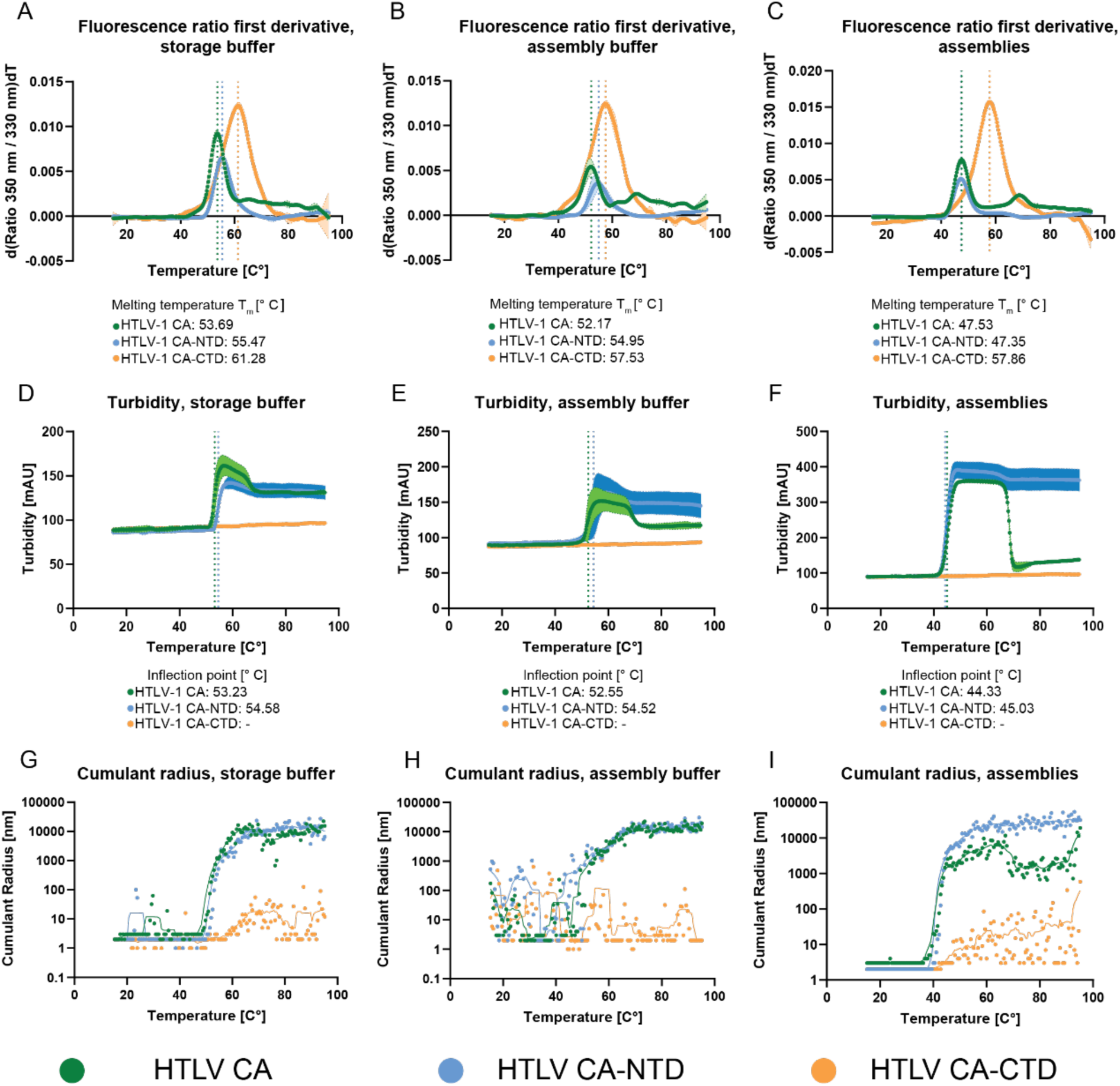
Increased assembly and aggregation propensity of CA-NTD. **A-C)** First derivative of the fluorescence ratio over a temperature ramp, measured by nanoDSF, for HTLV-1 CA (green), CA-NTD (cyan), and CA-CTD (orange) in **A)** non-assembled protein in storage buffer, **B)** non-assembled protein in assembly buffer, and **C)** protein in assembly buffer after assembly reaction. The dotted lines show the melting temperatures (Tm) of each construct, which correspond to unfolding events. **D-F)** Turbidity change as a function of temperature, measured by back-reflection, for HTLV-1 CA, CA-NTD, and CA-CTD in **D)** non-assembled protein in storage buffer, **E)** non-assembled protein in assembly buffer, and **F)** protein in assembly buffer after assembly reaction. The turbidity increases drastically for CA and CA-NTD in all conditions, but not for CA-CTD. The inflection points for the turbidity increase of CA and CA-NTD coincide with their melting temperatures (A-C). At higher temperature, CA turbidity levels decrease again markedly in both assembly conditions, and the inflection point for this decrease coincides with the second unfolding event of CA. **G-I)** Cumulant radius over a temperature ramp, measured by DLS, for HTLV-1 CA, CA-NTD, and CA-CTD in **G)** non-assembled protein in storage buffer, **H)** non-assembled protein in assembly buffer, and **I)** protein in assembly buffer after assembly reaction. CA and CA-NTD show an increase in cumulant radius coinciding with the observed turbidity increase and respective melting temperatures. CA-CTD shows no significant cumulant radius increase across all conditions. The color code is indicated.

This data revealed overall similar behavior of full CA and CA-NTD, while CA-CTD behaved differently. CA-CTD reproducibly showed the highest melting temperature (Tm) in all three conditions, suggesting that CA-CTD is the most stable of the three protein variants we tested. Moreover, full-length CA showed two unfolding events when assembled. Notably, CA-NTD shifted to a lower melting temperature when assembled, matching more closely to the temperature of the first unfolding event for full-length CA. This suggests that CA-NTD is the first domain of CA to unfold. Both full-length CA and CA-NTD showed a higher propensity to either aggregate or potentially form regular structures (**see Figure 3D-I**), while CA-CTD did not show an increase in turbidity or cumulant radius in any of the conditions tested.

Taken together, these measurements are consistent with our structural observations and further confirm the assembly-defining role of the CA-NTD, where the aggregation/assembly propensity in full-length CA is clearly derived from the CA-NTD and not the CA-CTD.

## Discussion

### A non-canonical stabilization of immature HTLV-1 by the CA-NTD

The work presented here expands our knowledge of the immature virion Gag architecture to five of the seven genera within the *Retroviridae* family (**Figure S3**). Importantly, using cryo-ET and subtomogram averaging of cell-derived VLPs, we identified a novel and unconventional mechanism of immature Gag lattice stabilization that is, to date, unique to HTLV-1. Our structural and biophysical experiments are consistent with a mode in which the CA-NTD is the only lattice-forming CA domain in immature HTLV-1 particles. This is supported by a previous study that did not find mutations in the CA-CTD affecting particle budding (*28*) and that in our *in vitro* MA126CANC assemblies the CA-CTD is less organized than the CA-NTD (or the CA-CTD of other retroviruses).

The structural interactions that differentiate retroviral genera can be defined as contributions of the individual interfaces and the orientation of the CA-NTD with respect to the local symmetry axes. The CA-NTD arrangement in HTLV-1 (*Deltaretroviru*s) is similar to that of MLV (*Gammaretrovirus*), with both having a tight packing of the trimerization interface, and helix 1 pointing towards the center of the hexamer. This results in positioning of CA-NTD in HTLV-1 and MLV closer to that observed in mature CA lattice conformations (*18*).

### The differential roles of HTLV-1 CA-NTD and CA-CTD in immature lattice stabilization

A unique feature reported here for HTLV-1 is the varying distance of the CA layer to the viral membrane in immature Gag-based VLPs. These differences in spacing correspond to the differences in curvature and flat patches of the HTLV-1 Gag lattice as reported here (**Figure 1A**) and previously (*21*).

Unlike RSV and MPMV, which also display a larger, but uniform distance of their CA layer to the viral membrane (*8*, *39*), HTLV-1 does not contain any non-canonical Gag domains between MA and CA. Therefore, the distance between membrane and CA layer is likely determined by specific properties of MA and CA and not the presence of additional domains.

A study of HIV-1/HTLV-1 Gag chimeras, showed that the HTLV-1 CA-NTD is the curvature-defining element, while MA regulates the CA-membrane distance (*40*). This was convincingly demonstrated, as any construct with the CA-NTD from HTLV-1 contained a flat lattice. A curved membrane over a flat lattice was observed only in those constructs where both CA-NTD and MA from HTLV-1 were present. The clear preference of HTLV-1 CA-NTD to shape lattices into flat regions appears to be compensated for by more flexibility in HTLV-1 Gag layers both above and below it. On the proximal side, MA seems to tolerate a larger membrane-CA distance range (**Figure S2A**). On the distal side, CA-CTD forms sparser lateral interactions than other retroviruses, which makes the location of CA-CTD dimers less constrained with respect to the CA-NTD network. This is apparent from the blurring of the CA-CTD layer in our subtomogram averages. Accordingly, residues in the MHR, which have been implicated in immature assembly for most retroviruses, are not positioned to form interactions stabilizing the immature HTLV-1 CA-CTD hexamer. To date most retroviruses have a structural stabilization element, which forms at the 6-fold symmetry axis below the CTD of CA (*9*, *18*, *25*, *35*, *41*, *42*). Our results here find no such domain in HTLV-1, further highlighting that HTLV-1 is structurally unique.

The fundamental difference in the organization of the CA-CTD in HTLV-1 compared to other retroviruses is quite striking. While for other retroviruses the CA-CTD is a critical lattice-forming element (*6*, *23*, *36*, *43*), in HTLV-1 some of the canonical CA-CTD functions are taken up by the CA-NTD (**Figure 4**).

**Figure 4:**
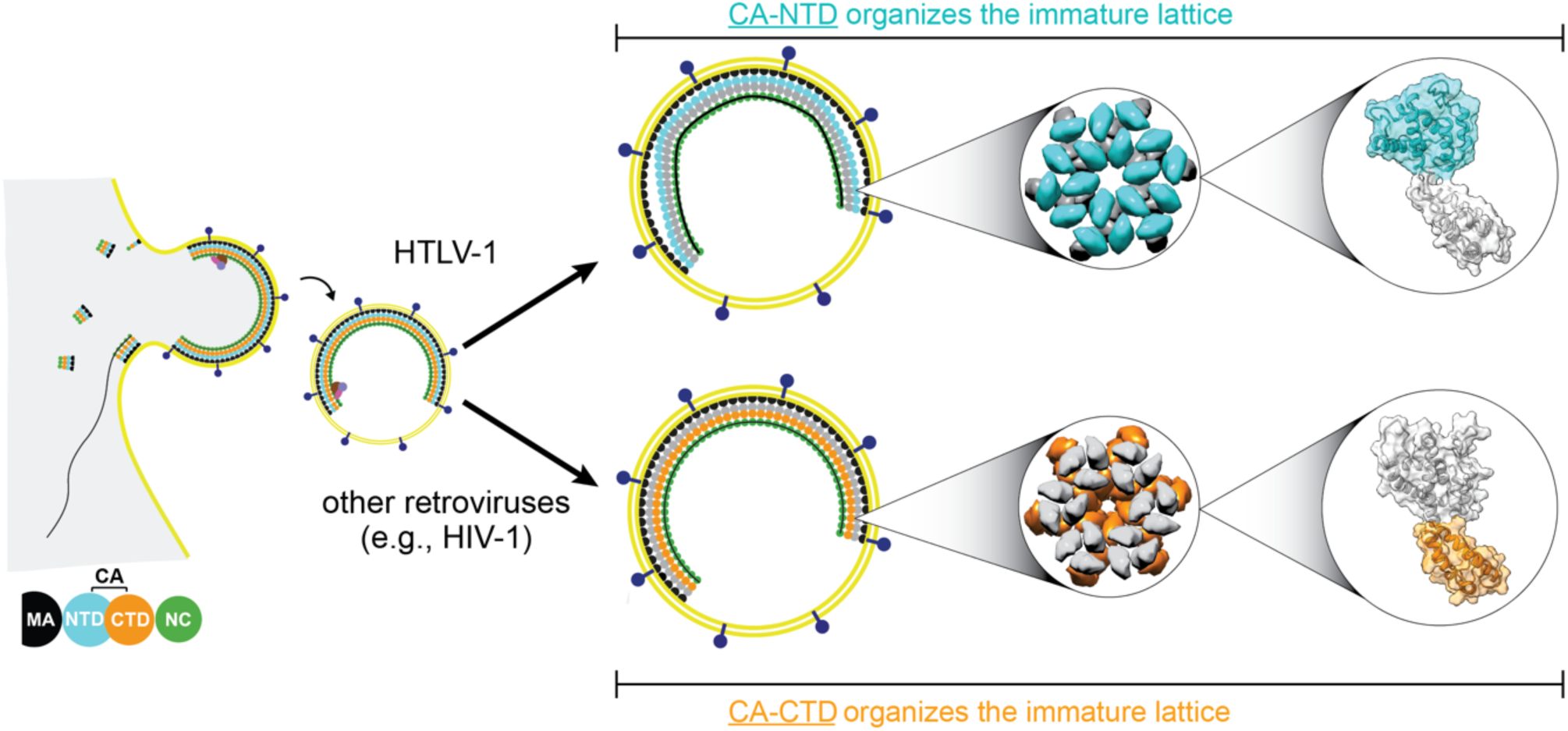
Unconventional immature HTLV-1 CA lattice stabilization. Schematic representation of the differential role of the CA-NTD and CA-CTD in the assembly of immature HTLV-1 particles, compared to other retroviruses with known immature structures (using HIV-1 as an example). In the case of HTLV-1, immature lattice interactions are driven by the CA-NTD, while in HIV-1, the CA-CTD is driving particle formation.

### The role of the HTLV-1 CA-CTD in immature HTLV-1 assembly

This raises the question; what is the function of the CA-CTD in immature HTLV-1 assembly? One possibility is that in HTLV-1 a direct interaction between the NC-RNA complex and the CA-CTD may contribute to immature lattice formation. In support of this hypothesis, HTLV-1 CA-CTD possesses a stretch of positively charged residues at its base (towards the VLP core) in helices 9 and 10 which could interact with NC-and/or-RNA (see underscored residues in **Figure S1B**). This would be similar to what was previously for M-PMV, where a basic RKK motif at the CA-CTD base is involved in promoting virus assembly and RNA packaging (*44*). EIAV also has a basic motif, RHR, and in slightly acidic *in vitro* assembly conditions, density is observed to interact with this motif, which is consistent with the proposed model (*26*).

Moreover, the C-terminal region of the NC domain of HTLV-1 Gag is negatively charged, a feature absent in other retroviruses. Accordingly, it has been shown that HTLV-1 NC is a weak RNA chaperone and nucleic acid binder (*45*). The authors pose the hypothesis that the NC C-terminus interacts intramolecularly with its’ zinc fingers effectively blocking non-specific RNA binding. It was further suggested that after Gag oligomerization into a lattice, lateral inter-molecular NC interactions form between the C-terminus and zinc fingers. Additional studies are warranted to determine if one of these models more accurately describes the interactions in this region of the immature lattice.

Our biophysical analyses are consistent with our structural observations that the key self-assembly/aggregation properties of CA primarily stem from the CA-NTD (**Figure 3D-I**). Interestingly, nanoDSF showed an additional melting temperature in CA under assembly conditions that is not present in either non-assembled CA or in CA-NTD under assembly conditions (**Figure 3A**). This indicates that the CA-CTD may further stabilize the lattice formed by the CA-NTD by expanding the interaction network through dimerization. Interestingly, a previous study used nanoDSF to study HIV-1 CA stability as a function of temperature, and suggested a reduced stability of the CA-CTD compared to the CA-NTD (*46*), in contrast to what we have reported here for HTLV-1. These differences in CA-CTD stability and interaction potential provide exciting avenues for further studies to better understand the differential retroviral CA assembly mechanisms.

## Conclusions

HTLV-1 infectious spread occurs predominantly by cell-to-cell contact and not by cell-free virus infection of permissive T-cells. The implications of this behavior for immature Gag lattice formation and maturation are unclear. Future work using cellular tomography on assembling and budding virions could provide a clearer understanding of the structural intermediates in the HTLV-1 lifecycle that are important in addressing this outstanding question in the field.

In addition to HTLV-1, the *Deltaretrovirus* genus contains multiple members, such as bovine leukemia virus. Comparing the assembly mechanisms among deltaretroviruses could help us better understand the general principles within this genus, and could clarify whether the observed dominance of the HTLV-1 CA-NTD in the immature Gag lattice is conserved within the genus. Our findings provide a structural basis for guiding future studies on HTLV-1 assembly and maturation, as well as for guiding the discovery of targets for therapeutic intervention. The distinct and unique nature of the HTLV-1 lattice, *i.e.,* the absence of hexameric CTD interactions and of a clear IP6 binding site as observed in HIV-1 (*47*), will necessitate alternative drug targeting approaches than, for example, the structure-based maturation inhibitors developed against HIV-1 (*48–50*).

## Acknowledgements

This work was funded by the Institute of Science and Technology Austria (ISTA) and the Austrian Science Fund (FWF) grant P31445 to F.K.M.S. Access to high-resolution cryo-ET data acquisition at EMBL Heidelberg was supported through the EMBL cryo-EM platform. We thank Victor-Valentin Hodirnau at ISTA, and Wim Hagen and Felix Weis at EMBL Heidelberg for support in cryo-ET data acquisition. This research was also supported by the Scientific Service Units (SSUs) of ISTA through resources provided by Scientific Computing (SciComp), the Life Science Facility (LSF), and the Electron Microscopy Facility (EMF). L.M.M. was supported by NIH grants R01 GM151775 and R21 DE032878 and by the University of Minnesota Masonic Cancer Center.

R.A.D was supported by National Institute of Allergy and Infectious Diseases (NIAID) grant R01AI147890. Specifically, we want to thank Alois Schlögl for computational support and Jesse Hansen and Volker Vogt for critical comments on the manuscript. We also thank the other members of the Schur lab for helpful discussions and experimental advice.

## Data availability

The electron microscopy density maps and models for the immature HTLV-1 CA-NTD lattice and representative tomograms have been deposited in the Electron Microscopy Data Bank (accession codes: EMD-17929, EMD-17930, EMD-17931, EMD-17932, EMD-17933, EMD-17934, EMD-17935, EMD-17936, EMD-17937, EMD-17938, EMD-17939, EMD-17940, EMD-17941, EMD17942, EMD-17943) and the Protein Data Bank (accession codes: 8PU6, 8PU7, 8PU8, 8PU9, 8PUA, 8PUB, 8PUC, 8PUD, 8PUE, 8PUF, 8PUG, 8PUH).

## Author contributions

Project administration: F.K.M.S.; supervision and funding acquisition: L.M.M, R.A.D, F.K.M.S.; conceptualization: M.O. and F.K.M.S.; methodology: M.O, and F.K.M.S.; investigation: M.O, M.P, D.C, H.Y., A.T., G.P, D.P, R.A.D, F.K.M.S; Software: M.O.; validation, formal analysis, and visualization: M.O., M.P, H.Y, F.K.M.S.; data curation: M.O. and F.K.M.S.; writing—original draft: M.O and F.K.M.S.; writing—review and editing: M.O., M.P, D.C, H.Y, A.T, G.P, D.P. L.M.M, R.A.D and F.K.M.S

## Materials and Methods

### Mammalian cell culture and Gag-based VLP production and purification

Codon optimized HTLV-1 Gag sequence, subcloned into the pCMV expression vector was ordered from Thermo Fisher Scientific. 293T cells, cultured in 10 x 10cm dishes, were transfected with the expression plasmid using Lipofectamine LTX Reagent with PLUS Reagent (Thermo Fisher Scientific, no. 15338100) according to manufacturer’s instructions. Cultivation medium, containing released HTLV-1 Gag-based VLPs was harvested 40 hours post transfection by ultracentrifugation. This and all further centrifugation steps were performed at 4°C. Cultivation medium was first clarified by centrifugation for 5 min at 1500 x *g* and subsequent filtering through syringe mounted PVDF filter with 0.45 µm pore size (Merck Millipore). VLPs were pelleted through a 20% sucrose cushion at 125,000 x *g* for 120 min. The pellet was briefly air-dried and resuspended in 150 μl PBS. Pooled and resuspended pellets were applied on a 6-18% Optiprep gradient, and centrifuged at 235,000 g for 90 min. Fractions containing Gag-based VLPs were pooled, diluted 1:4 in PBS, and pelleted without a cushion at 260,000 x *g*, for 45 min. The final pellet was then resuspended in 12 μl of PBS.

### Cryo-electron tomography for HTLV-1 Gag-based VLPs

#### Grid preparation for HTLV-1 Gag-based VLPs

The resuspended Gag-based VLP pellet was kept on ice and mixed with 10 nm colloidal gold suspension prior to vitrification. 2.5 µl of solution was applied on glow-discharged C-flat 2/2 3C grids and vitrified in liquid ethane using a Leica EM GP2 plunge freezer. Grids were blotted in back side mode for 3-4s. The humidity chamber was conditioned to 8°C and 90% relative humidity. Grids were stored in liquid nitrogen until imaging.

#### Imaging of HTLV-1 Gag-based VLPs

Data were collected on a Thermo Fisher Scientific Krios G3i equipped with Gatan Quantum K3 using SerialEM for data acquisition (*51*). Areas of interest for high-resolution data collection were identified in low magnification montages. Prior to tomogram acquisition, gain references were acquired and the filter was fully tuned. Microscope tuning was performed using the FEI AutoCTF software or SerialEM tuning functions. The slit width of the filter was set to 20 eV. The magnification was set to 80,000x, resulting in a pixel spacing of 1.381 Å. Tilt images were acquired as movies in super-resolution mode consisting of 8 dose-fractions. The dose rate was set at ∼18 electron/pixel/second. Tilt series were acquired using a dose-symmetric tilt-scheme (*52*), with a tilt range from 0° to −60° and +60° in 3° steps, and at nominal defocus between -1.25 and -3.5 µm. The total dose per tilt image was 3.5 e-/Å^2^.

#### Cryo-ET image processing of HTLV-1 Gag-based VLPs

A schematic overview over the individual processing steps for the HTLV-1 Gag-based VLPs is shown in **FigureS2C**. Tilt images with blocked field of view were removed prior to further steps. Tilt series were aligned using the Etomo package and exposure-filtered, as published previously, using a custom Matlab script (*9*, *53*, *54*).

Template matching was done essentially as previously published (*55*). A template was generated using subtomogram alignment of the CA layer from round VLPs from 12 tomograms. A cylindrical mask centered at a CA hexamer and encompassing two adjacent CA rings was applied to the template. In-plane and cone angles were scanned in steps over a range of 180° and 360°, in 6° and 8° steps, respectively. Lattice connectivity analysis (as described in (*55*)) was performed to remove peaks outside of lattice, and annotate VLPs. The following criteria were used to assess pairs of local cross-correlation maxima: minimum/maximum spacing between hexamers was 45 Å/110 Å, minimum/maximum curvature was= −15°/25°. Only networks containing >20 cross-correlation peaks were considered as patches of lattice and used for subsequent steps.

For initial subtomogram alignment, tomograms were reconstructed in novaCTF (*56*) using 2D CTF correction of 2x binned stacks with the multiplication algorithm. Bin8 and bin4 alignments were performed in Dynamo (*57*) and subTOM (*26*). A conservative lowpass filter of ≥25 Å was used during these alignments. Given the irregular and variable shape of Gag-based VLPs, we refrained from using C6 rotational symmetry, which is conventionally used for processing of immature retroviral lattices. Instead, we used C2, which allows for capturing CA hexamer deformation. After bin4 alignments the VLP list was manually curated to remove particle formations outside enveloped VLPs.

Preprocessing for the Warp-Relion-M pipeline was done in Warp (*58*). Tilt series alignments from Etomo and subtomogram alignments from Dynamo/subTOM were imported and used for subtomogram reconstruction. 3D refinement was performed in Relion 3.0.8 at bin4 and bin2, using subtomograms reconstructed in Warp at the respective binning. 3D classification was performed in Relion 3.0.8 (*59*) at bin2 to select a more homogeneous CA-CTD population. A cylindrical mask encompassing six CA-CTD dimers was used during the classification. For both CA-NTD and CA-CTD, a separate species was set up masking the respective layer. The CA-NTD species contained all particles from the 3D refinement at bin2, whereas the CA-CTD species contained just particles from a selected class (**Figure S2A**). Five rounds of multi-particle refinement were performed, refining particle positions, 2D and 3D warping, and CTF parameters. Frame alignment refinement was performed in the last two rounds.

#### Model building for the immature HTLV-1 CA lattice from the Gag-based VLP-derived structure

The model of the immature HTLV-CA assembly was generated via rigid body fitting the HTLV-1 CA-NTD derived from pdb 1QRJ (*12*) into our electron microscopy density using the fit in map option in UCSF Chimera. As we noted that the experimentally-derived HTLV-1 CA-CTD from pdb 1QRJ did result in severe clashes on the CA-CTD dimer interface upon rigid body fitting, we employed a computationally predicted model of the CA-CTD (CA residues 129-207) using ColabFold (*60*). Interestingly, this predicted model fitted into our EM-density better and resulted in less clashes across the dimeric interface. Hence, we used this rigid body fitted prediction for modelling the CA-CTD.

### Protein purification and *in vitro* assembly

#### Cloning of truncated Gag constructs for *in vitro* assembly

The sequences encoding MA126CANC, CA or just the CA-NTD and CA-CTD domains of HTLV-1 Gag were cloned into pET28 expression vector in frame with 6xHis-SUMO tags using standard molecular cloning methods.

#### Protein expression and purification of MA126CANC

An overnight culture of BL21 carrying the pET28/His-SUMO-MA126CANC construct was prepared to inoculate a total of 2 liters LB medium supplemented with Kanamycin. Following incubation at 37°C with shaking at 210 rpm until the bacterial culture reached OD600 of 0.5 – 0.7 protein expression was induced by addition of IPTG at a final concentration of 1mM. Induction was carried out for 6 hours at 37°C with shaking at 210 rpm.

Bacterial cells were harvested by centrifugation at 6,000 x *g* for 15 min. The resulting cell pellet was dissolved in resuspension buffer (20 mM Tris, 500 mM NaCl, 2 µM ZnCl2, 5% Glycerol, 1mM PMSF, 1 mM TCEP, pH 8.0)

Protein was extracted from cells by cell lysis *via* 3 cycles of freeze-thaw involving freezing at −80°C followed by thawing at 42°C. After 45-60 min of centrifugation of the lysate at 21,000 x *g* and 4°C, the supernatant was collected and subjected to nucleic acid precipitation by treatment with 10% PEI (Polyethylenimine) at a final concentration of 0.3% under stirring at 4°C for 30 min. Afterwards the mixture was centrifuged for 10 min at 10,000 x *g* and 4°C. The remaining supernatant was treated with ammonium sulfate at a final concentration of 40% under stirring at 4°C overnight for precipitation of His-SUMO-MA126CANC. Precipitated protein was collected by centrifugation at 10,000 x *g* and 4°C for 20 min.

Purification of His-SUMO-MA126CANC was carried out utilizing anion exchange chromatography and affinity chromatography. For anion exchange chromatography the protein was dissolved in buffer composed of 20 mM Tris, 2 mM TCEP at pH 8.0 and was then applied to a 5 ml HiTrap^TM^ SP HP column (Cytiva, Cat. No. 17115201) before being eluted with high salt buffer composed of 20 mM Tris, 500 mM NaCl, 2 mM TCEP at pH 8.0.

The protein sample was then transferred to a 1 ml HisTrap^TM^FF column (Cytiva, Cat. No. 17531901) for protein purification by affinity chromatography. Bound protein was treated with wash buffer (20 mM Tris, 500 mM NaCl, 20 mM Imidazole, 2 mM TCEP, pH 8.0) before it was eluted with high concentration Imidazole buffer (20 mM Tris, 500 mM NaCl, 250 mM Imidazole, 2 mM TCEP, pH 8.0).

To remove Imidazole, the protein solution was dialyzed overnight at 4°C in tubes of the Pur-A-Lyzer^TM^ Maxi 6000 Dialysis Kit (Sigma, cat. No. PURX60100-1KT) against dialysis buffer (20 mM Tris, 500 mM NaCl, 0.5 mM TCEP, pH 8.0). During dialysis the sample was treated with N-terminally His-tagged Ulp1 protease for removal of the His-SUMO tag from MA126CANC. The protease was later removed by reapplication of the sample to a 1 ml HisTrap^TM^FF column resulting in high affinity binding of His-Ulp1 to the Nickel sepharose resin. MA126CANC protein exhibits low affinity for the packing material of this column. Hence, it was eluted with 20 mm Imidazole wash buffer. After another dialysis step, 150 – 200 µl aliquots of purified protein were flash frozen in liquid nitrogen and stored at −80°C.

#### Assembly of MA126CANC tubes

Protein stored in dialysis buffer of 500 mM NaCl was first mixed with equal volume of salt-free buffer (20 mM Tris, 0.5 mM TCEP, pH 8.0). The protein sample was then transferred into a Pierce concentrator tube 10 kDa molecular weight cut off (MWCO) (Thermo Fisher Scientific, Cat. no. 88513) and protein concentration was increased to 12 – 20 mg/ml by centrifugation at 15,000 x *g* and 4°C for 12 min. 10 µl of protein (120 – 200 µg) were then mixed with 2 µl of 5 mg/ml GT50 nucleotides, 1 µl 50 mM EDTA and 37 µl of salt-free buffer.

The reaction mixture with a final NaCl concentration of 50 mM was incubated overnight at 4°C. After incubation an aliquot of the sample was subjected to negative staining for confirmation of VLP presence by transmission electron microscopy on a Tecnai T10.

#### Protein expression and purification of CA, CA-NTD and CA-CTD

E. coli in glycerol stocks with pET28a vectors containing a His-SUMO tag, with HTLV-1 CA, CA-NTD, and CA-CTD inserts respectively, were used to inoculate 50 ml of LB medium and grown overnight at 37 °C and 220 rpm until an OD600 of 2.5 was reached. Half of each overnight culture was used to inoculate 1L of LB medium and grown at 37°C and 220 rpm until an OD600 of 0.6 was reached, after which protein expression was induced with 1 mM IPTG for 6h. The cultures were centrifuged at 6,000 × *g* and 4°C for 15 min to collect the cell pellets which were subsequently dissolved in lysis buffer (20 mM Tris, 500 mM NaCl, 2 µM ZnCl2, 5% Glycerol, 1mM PMSF, 1 mM TCEP, pH 8.0) and subjected to 3-5 freeze thaw cycles to disrupt the cells.

The resulting lysed cells were centrifuged at 13,000 × *g* for 45 min to separate protein from insoluble cell debris, followed by protein purification from nucleic acid by precipitating the nucleic acid with 10% PEI at a final concentration of 0.3% for 10 min under stirring at 4°C.

The solutions were centrifuged again, at 10,000 × *g* and 4°C for 10 min to collect the precipitated nucleic acid and ammonium sulfate was subsequently added to the supernatant to a saturation of 40% under stirring at 4°C overnight to precipitate the proteins. The solutions were centrifuged again at 13,000 × *g* to collect the proteins in a pellet before resuspending the protein pellets in wash buffer (20 mM Tris, 20 mM imidazole, 500 mM NaCl, 2 mM TCEP, pH 8).

The proteins were then purified using affinity chromatography with 1 mL HisTrap^TM^ FF according to the manufacturers’ instructions (HisTrap FF columns, Cytiva) and elution buffer containing 20 mM Tris, 250 mM imidazole, 500 mM NaCl, 2 mM TCEP, pH 8.

Following purification, the His-SUMO tag was cleaved from the proteins using Ulp1 protease and the proteins placed for dialysis at 4°C overnight in dialysis buffer (20 mM Tris, 500 mM NaCl, 5 mM TCEP, and pH 8). A second affinity chromatography purification step with Ni-NTA columns was done to purify the proteins from the cleaved His-SUMO tag, followed again by dialysis in dialysis buffer at 4°C overnight.

#### Assembly reaction and preparation for nano differential scanning fluorimetry

HTLV-1 CA, CA-NTD, and CA-CTD were separately concentrated in Pierce concentrator tubes (10 MWCO for CA and CA-NTD, and 3 MWCO for CA-CTD). Monomeric HTLV-1 CA, CA-NTD, and CA-CTD were diluted to 1 mg/mL concentration in both dialysis buffer and under assembly conditions (50 mM MES, 200 mM NaCl, pH6) in a volume of 10 μl and placed in Prometheus Series High Sensitivity Capillaries (NanoTemper, Cat. no. PR-C006).

For measurement of assemblies the proteins were first placed in assembly buffer (50 mM MES, 200 mM NaCl, pH6) with a final protein concentration of 16 mg/mL and were incubated at 4°C for 4h and 26 °C overnight. Again, a volume of 10 μl was then placed in Prometheus Series High Sensitivity Capillaries (NanoTemper, Cat. no. PR-C006).

#### Nano differential scanning fluorimetry, backreflection, and static light scattering

The capillaries were loaded into a Prometheus Panta instrument (NanoTemper Technologies). NanoDSF, backreflection, and DLS were measured continuously over a temperature ramp of 5°C/min from 15°C to 95°C at 40% LED excitation power. All scans were single reads of two replicates performed at least three times.

### Cryo-electron tomography for HTLV-1 MA126CANC tubes

#### Grid preparation MA126CANC tubes

After *in vitro* assembly MA126CANC tubes were kept at 4°C until plunge freezing. 2/2-3C C-flat grids coated with 2 nm support carbon layer were glow discharged in the presence of amylamine. Grids were then incubated on a 5 μl drop of sample for 10 min. The rest of the sample was mixed with 10nm colloidal gold, and 2.5 μl of this solution was added to the incubated grids prior vitrification. The samples were vitrified in liquid ethane using a Leica GP2 plunger with front-side blotting (blot time of 3-4 s, humidity 90-95%, temperature 10°C) and stored in liquid nitrogen until imaging.

#### Imaging of MA126CANC tubes

For *in vitro* assembled MA126CANC tubes we acquired data on two systems. One dataset was collected on a Thermo Fisher Scientific Krios G3i equipped with Gatan Quantum K3 (system 1). The second dataset was collected on a FEI Titan Krios, operated at 300 keV, equipped with a Gatan Quantum 967 LS energy filter and a Gatan K2xp direct electron detector (system 2). The slit width of the filter was set to 20 eV on both systems. SerialEM was used for data acquisition in both cases. Areas of interest for high-resolution data collection were identified in low magnification montages. Prior to tomogram acquisition, gain references were acquired and the filter was fully tuned. Microscope tuning was performed using the FEI AutoCTF software or SerialEM tuning functions.

On system 1 the magnification was set to 80,000x, resulting in a pixel spacing of 1.381 Å. Tilt images were acquired as movies in super-resolution consisting of 8 dose-fractions. The dose rate was set at ∼21 eps. On system 2 the magnification was set to 105,000x, resulting in a pixel spacing of 1.327 Å. Tilt images were acquired as movies in super-resolution consisting of 1o dose-fractions. The dose rate was set at ∼ 5.4 eps.

Tilt series were acquired using a dose-symmetric tilt-scheme (*52*), with a tilt range from 0° to −60° and +60° in 3° steps, at nominal defocus between -1.5 and -3.5 µm. The total dose per tilt image was 3.5 e-/Å^2^.

#### Cryo-ET image processing of MA126CANC tubes

A schematic overview over the individual processing steps for the MA126CANC tubes is shown in **Figure S6A**. Prior to tilt series alignment, movies were aligned using the IMOD alignframes function. Tilt images with blocked field of view were removed prior to further steps. Tilt series were aligned using the Etomo package and exposure-filtered using a custom Matlab script (*9*, *53*). For initial subtomogram alignment (see down below), CTF-corrected tomograms were reconstructed in novaCTF using the multiplication algorithm with the slab size of 15nm. Bin8 and bin4 alignments were performed in Dynamo/Subtom. A conservative lowpass filter of ≥25 Å was used during these alignments.

Preprocessing for the Warp-Relion-M pipeline was done in Warp. Tilt series alignments from Etomo and subtomogram alignments from Dynamo/Subtom were imported and used for subtomogram reconstruction. 3D refinement was performed in Relion 3.0.8 at bin4 and bin2, using subtomograms reconstructed in Warp at the respective binning.

#### Classification of tube geometries for structure determination

Since the tube geometry was variable for the CANC tubes, tubes were sorted into groups containing similar tube geometries. Specifically, the angle between the tube axis and the hexamer-hexamer connection was used to distinguish between different tube geometries (**Figure S6B**). To this end, first a *de novo* reference was generated for each tube separately by subtomogram alignment of regularly spaced volumes along a given tube using Dynamo/Subtom. The initial alignments were performed at bin8. The first step was to remove particles that converged to the same spatial position using distance cleaning. A cross-correlation threshold was set manually for each tomogram upon visual inspection, to remove misaligned and bad subvolumes. The positions of subtomograms derived from the initial alignments were used for geometry analysis. Tubes were sorted into groups so that the maximum angle difference within one group was 5°. Subtomogram alignment and tube grouping was then refined at bin4.

#### Merging datasets for multiparticle refinement

Since the datasets acquired on the two microscope systems differed in pixel size, they were processed separately until the multiparticle refinement step. Then, since pixel size of system 1 was not calibrated, we used the fitmap function of UCSF Chimera (*61*) to maximize the overlap between maps generated using system 1 and 2, while varying pixel size of system 1. For further processing we used this estimated pixel size rather than the nominal value. Based on this, we estimated the actual pixel size on system 1 to be 1.339 Å. In order to unify the pixel size used for Multiparticle refinement, we used a box size which yields an integer upon dividing with both pixel sizes. A box size of 321.3 Å was fulfilling the criteria.

#### Multispecies refinement of MA126CANC tubes

3D refinement in Relion 3.0.8 was performed for each group of tubes separately at bin4 and bin2. Subtomograms were reconstructed in Warp. The respective halfmaps and particle coordinate star files were used as a starting point for multi-particle refinement in M. Five rounds of refinement were performed, gradually adding refinement options similarly as described in (*62*). This yielded nine CA-NTD hexamer structures at resolutions from 3.7 to 7.0 Å. In order to increase the resolution for model building, the two most populated groups sharing similar geometry were pooled, subtomograms were reconstructed using M at a pixel size of 1.77 Å, and the pooled population was subjected to Autorefine 3D in Relion. Afterwards, two final rounds of M refinement were performed. The final 3D refinement was performed in Relion, with subtomograms reconstructed using M at bin1.

### Model refinement and data visualization

The initial model for HTLV-1 CA-NTD was obtained by adjusting pdb 1QRJ, via flexible fitting it into the EM-density using the Isolde plugin of ChimeraX (*63*, *64*). The final model was then built by iterating between real space refinement in Phenix (*65*) and manual adjustments in Coot (*66*).

### Data visualization and figure preparation

3D volumes were visualized using UCSF Chimera (*61*), ChimeraX (*63*, *64*) and IMOD (*54*). Images of 3D volumes and models were made using Chimera or ChimeraX or via ray-tracing in Pymol (*67*). Figures were prepared using Adobe Illustrator (Adobe Inc.). Movies were generated in ChimeraX and Adobe Premiere Pro 2023 (Adobe Inc.).

1D density plots were calculated from bin1 (CANC tubes) or bin2 (Gag VLPs) unsharpened unmasked average, assuming a tube diameter of 60 nm and VLP diameter of 130 nm.

### Mutagenesis of residues involved in stabilizing the immature CA-NTD

In order to characterize the residues involved in stabilizing the immature CA-NTD on particle production, we conducted mutagenesis on key residues. First, we created residue L43A, as L43 may contact M17 and Y61. Second, we conducted mutagenesis of the trimeric CA-NTD interface. Specifically, residue pairs H72/Q73, Y90/N91, P92/L93, G95/P96, L97/R98, and V99/Q100 in the CA-NTD were created by using the Gibson assembly method as previously described (*68*). Verification of the correct creation of the mutants was done by Sanger DNA sequencing analysis. The efficiency of immature particle production was analyzed by quantifying Gag proteins in released particles through harvesting cell culture supernatants and by using immunoblot analysis with a mouse monoclonal anti-HTLV p24 antibody (Catalogue #: sc-53891; Santa Cruz Biotechnology, Dallas, TX). To determine particle production (i.e., particle release) efficiency of mutants relative to that of WT, the Gag expression levels detected in cell lysates of the WT and mutants were normalized relative to the respective GAPDH level. Mutant particle release to that of WT was then determined by the ratio of Gag levels detected from cell culture supernatants to that from the normalized Gag levels from the cell lysate, with WT particle release being set to 100 and particle release of the mutants being relative to that of the WT. Results were plotted by using GraphPad Prism 6.0 (GraphPad Software, Inc., CA). Relative significance between the WT and a mutant was determined by using an unpaired t-test. Cryo-EM analysis of immature particle morphologies of the L43A mutant was done by producing VLPs from cells, and concentrating by ultracentrifugation.

## Supplementary Figures

**Figure S1:**
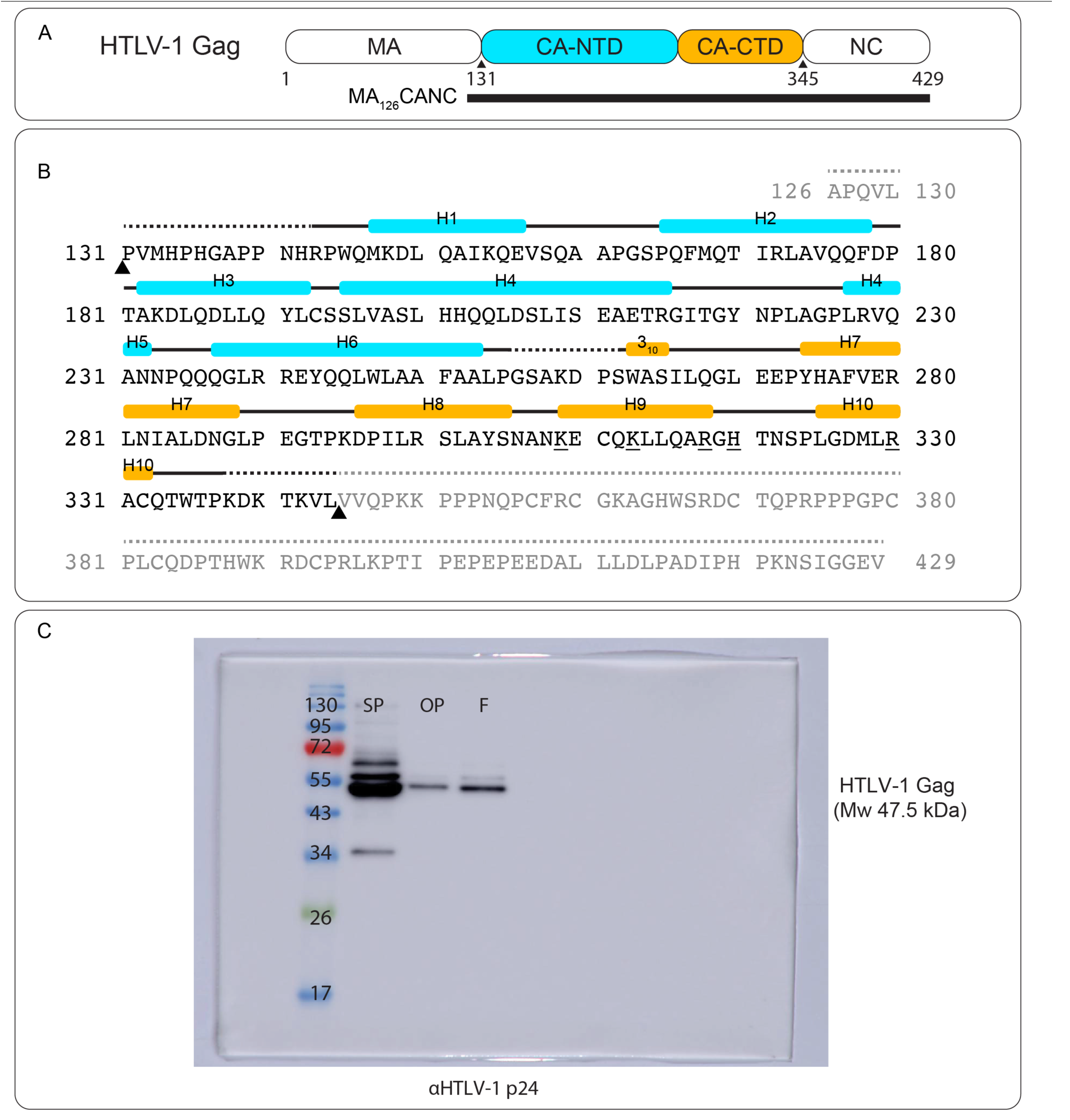
HTLV-1 Gag/CA sequence and Gag VLP purification. **A)** Schematic depiction of the domain architecture of HTLV-1 Gag and the MA126CANC truncation protein used for *in vitro* assembly and structure determination. Residue numbers and black triangles denote protease cleavage sites. Note the absence of non-canonical Gag domains in HTLV-1. **B)** Sequence of HTLV-1 MA126CANC. The positions of **α**-helices are shown as cyan and yellow bars. Regions for which structural data is lacking are dashed. Triangles denote protease cleavage sites. Please note that due to the used expression and purification system of our 6xHIS-SUMO construct, an ectopic Serine residue remains at the very N-terminus of the purified protein (not shown in this presented sequence). The underlined residues in Helix 9 and 10 denote positively charged residues which could form potential interactions with the NC-RNA. **C)** Samples of Optiprep gradient (OP)-purified particles were separated by SDS–polyacrylamide gel electrophoresis. Proteins were visualized by immunoblot using an anti-HTLV-1 p24 antibody. Sucrose pellet (SP), Optiprep bottom (OP), Final Opti prep pellet (F). Positions of molecular mass standards (in kiloDaltons, kDa) are indicated. The molecular weight (MW) of HTLV-1 Gag of 47.5 kDa is annotated.

**Figure S2:**
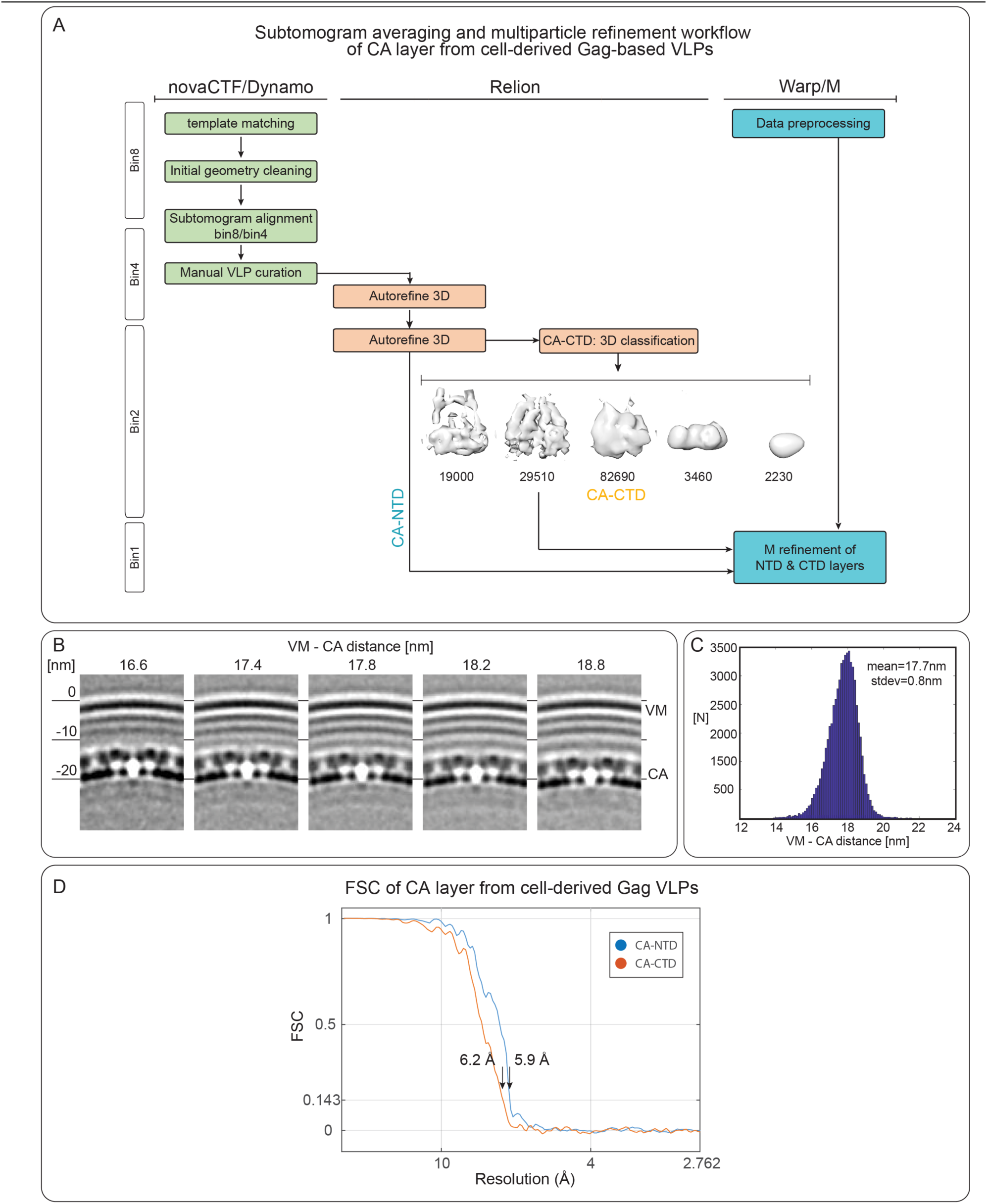
Image processing of HTLV-1 Gag VLPs. **A)** Image processing workflow for the subtomogram averaging and multiparticle refinement of the CA layer from HTLV-1 Gag-based VLPs. Kindly see Materials and Methods for more details. **B)** XZ-slices through bin4 averages of the immature HTLV-1 Gag lattice, showing the varying distance of the CA-layer with respect to the viral membrane (VM). The distance was measured from the outer VM layer to the electron-lucent layer between CA-NTD and CA-CTD. **C)** Histogram showing the measured distances of the CA-layer to the viral membrane. The distance was determined by re-aligning particles to viral membrane layers. The individual CA-membrane distances then correspond to the mean value + particle Z-shift. N=51,700. **D)** Fourier-shell correlation (FSC) for the independently aligned CA-NTD (blue line) and CA-CTD layer (red line), showing a resolution of 5.9 Å and 6.2 Å at the 0.143 criterion, respectively.

**Figure S3:**
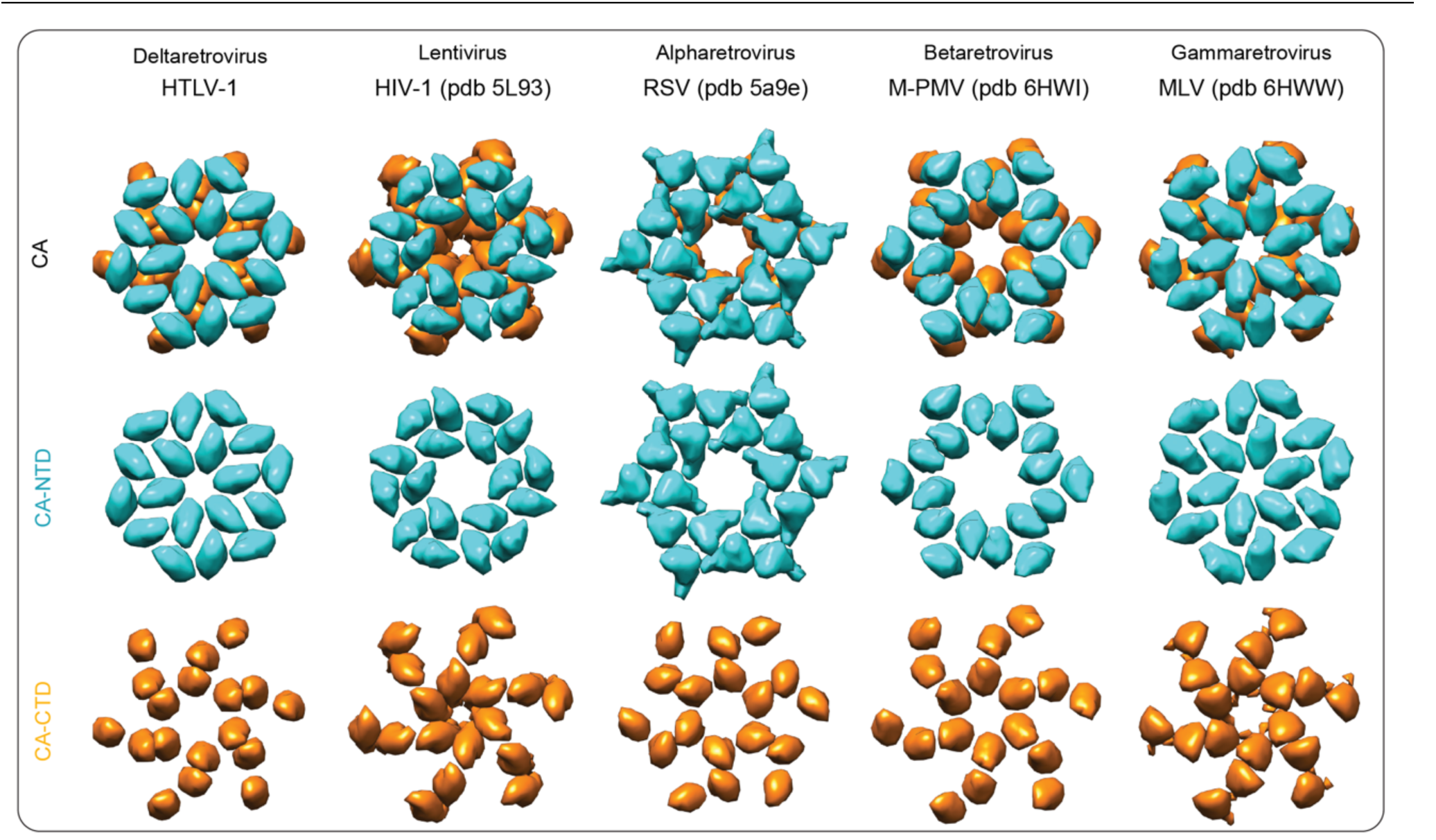
Immature quaternary CA arrangements among retrovirus genera. Schematic representation of the CA-NTD and CA-CTD arrangements within immature retroviruses from different genera. HTLV-1, *Deltaretrovirus* (as determined in this study); HIV-1, *Lentivirus*; RSV, *Alpharetrovirus*; M-PMV, *Betaretrovirus*; MLV, *Gammaretrovirus*.

**Figure S4:**
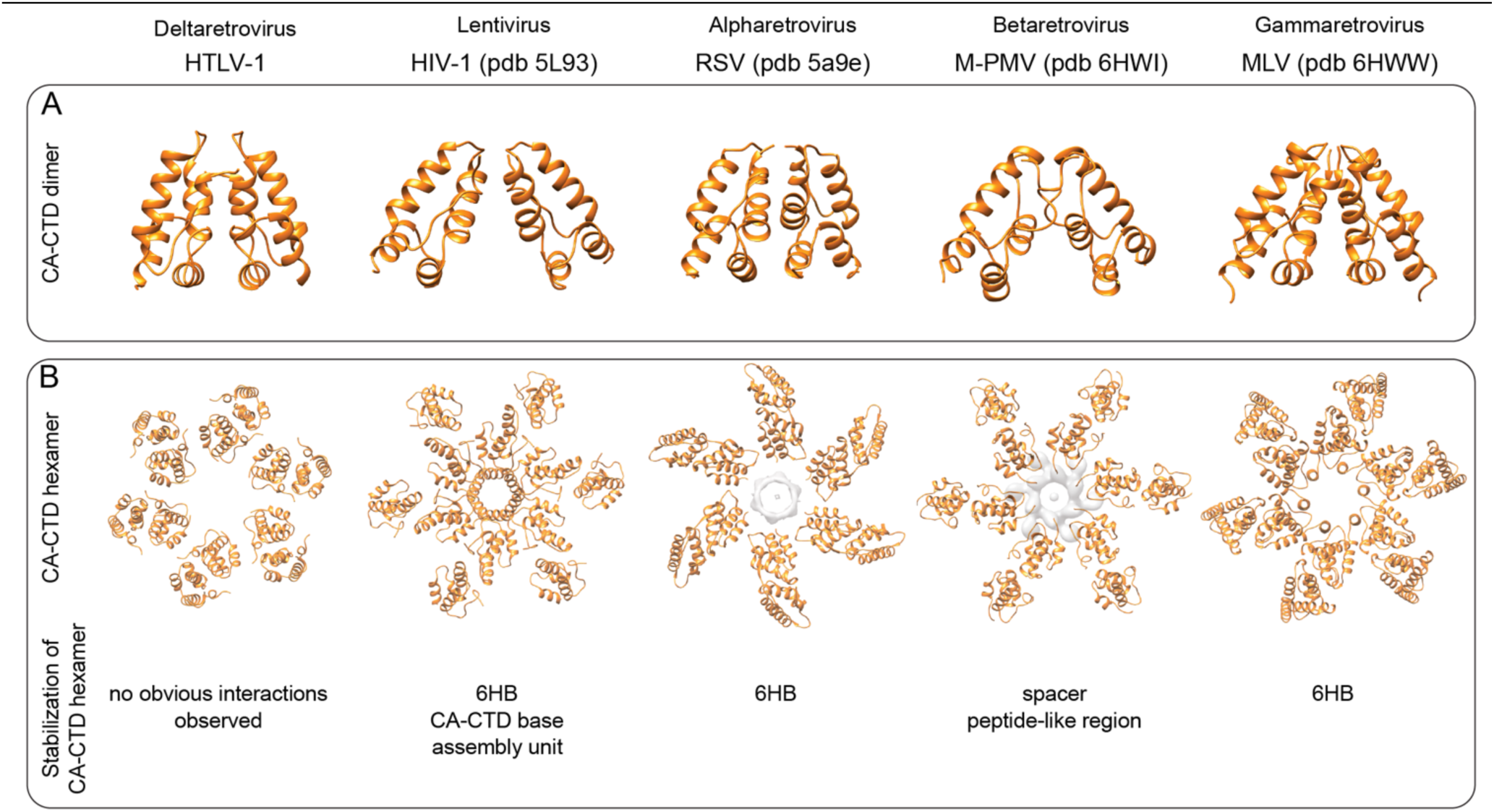
CA-CTD interactions in immature retroviral lattices. **A)** Comparison of different retroviral immature CA-CTD dimer structures. The pdb accession codes for the different retroviruses are indicated. The same models have been used to generate the data shown in panel (B). **B)** Comparison of the immature CA-CTD hexamer assembly. The main interaction interfaces described previously to stabilize the hexamer are annotated. The CA-CTD assembly unit in HIV-1 is described in (*9*). For RSV and M-PMV the electron microscopy densities for the RSV SP and the M-PMV spacer peptide-like region are shown, as for these regions no molecular model has yet been obtained.

**Figure S5:**
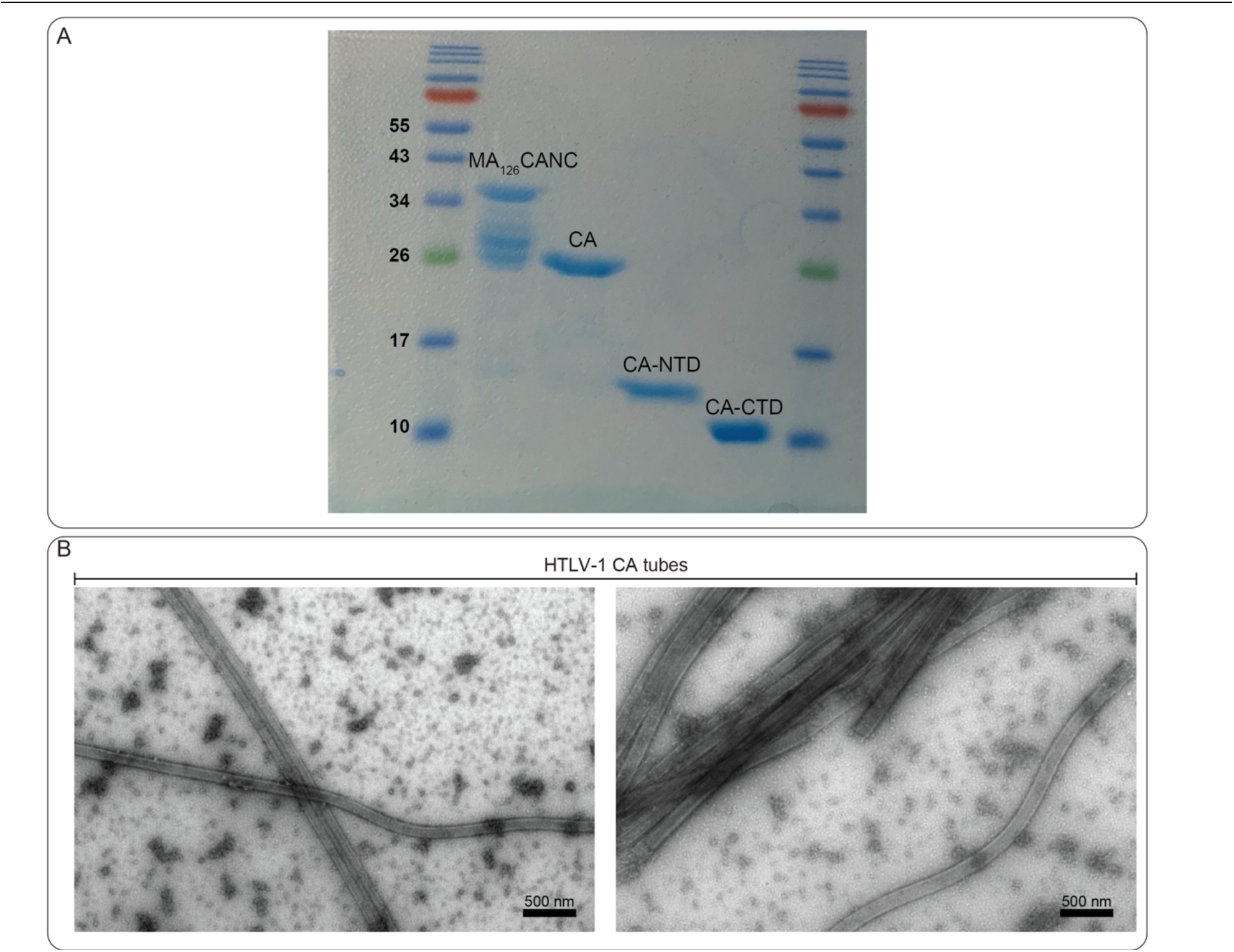
Expression and purification of proteins for *in vitro* experiments. **A)** Samples of purified CA protein variants were separated by SDS–PAGE (15%) and stained via Coomassie brilliant blue. Positions of molecular mass standards (in kiloDaltons, kDa) are indicated. **B)** Negative Stain TEM micrographs of HTLV-1 CA tubes. Scale bar is 500 nm.

**Figure S6:**
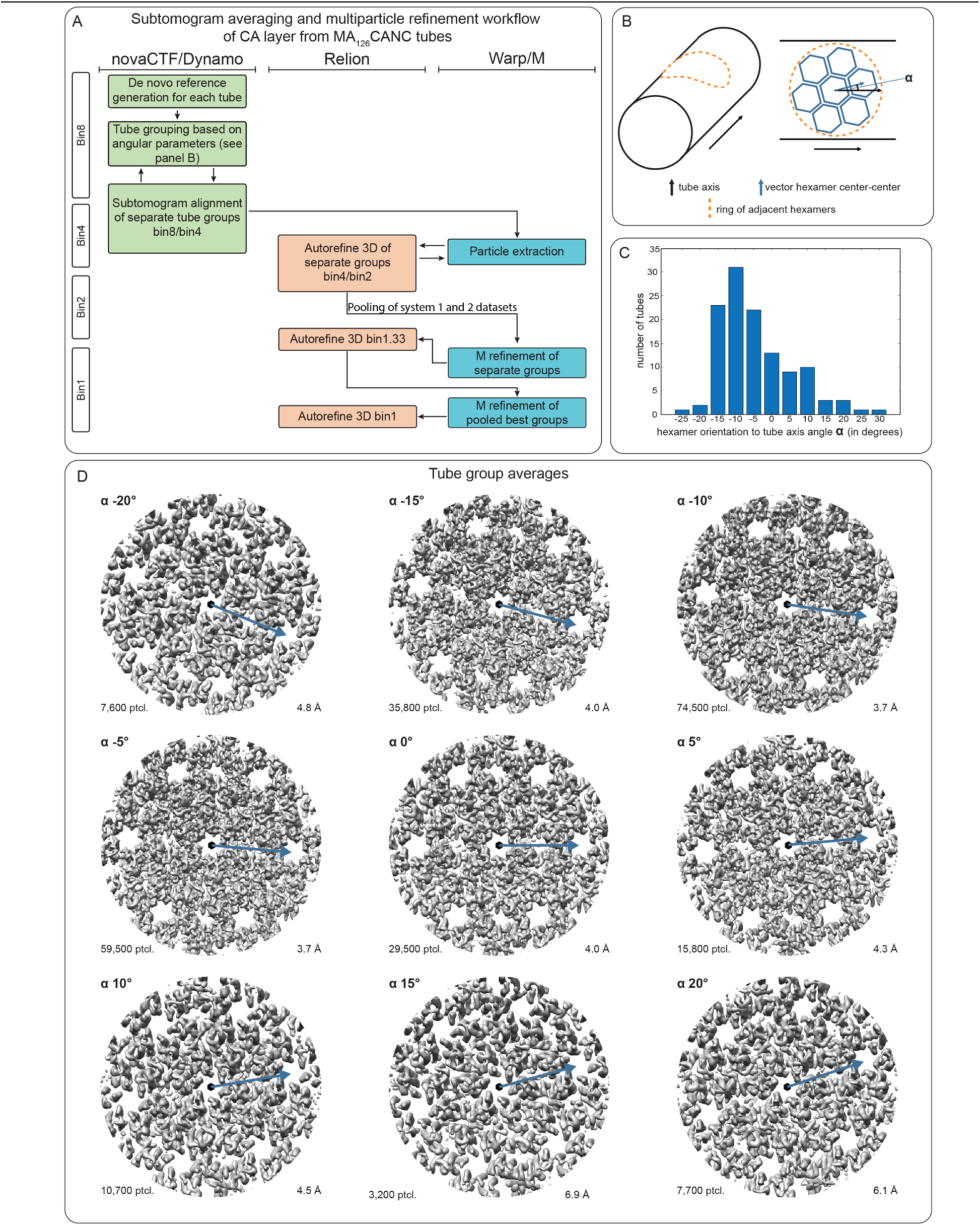
Image processing and classification of HTLV-1 MA126CANC tubes. **A)** Image processing workflow for the subtomogram averaging and multiparticle refinement of the CA-NTD layer from HTLV-1 MA126CANC tubes. Please see Materials and Methods for more details. **B)** Classification of tubes based on geometry. Different tubes have different geometries, with varying helical rise and pitch. For improved structure determination, tubes were grouped into classes according to their hexamer orientation (blue arrow) with respect to the tubes axis (black arrow). **C)** Histogram of abundance of different tube groups. **D)** Refined EM-density maps for the individual tube groups. Upper left corner – angle group; lower left corner – number of particles; lower right corner –resolution.

**Figure S7:**
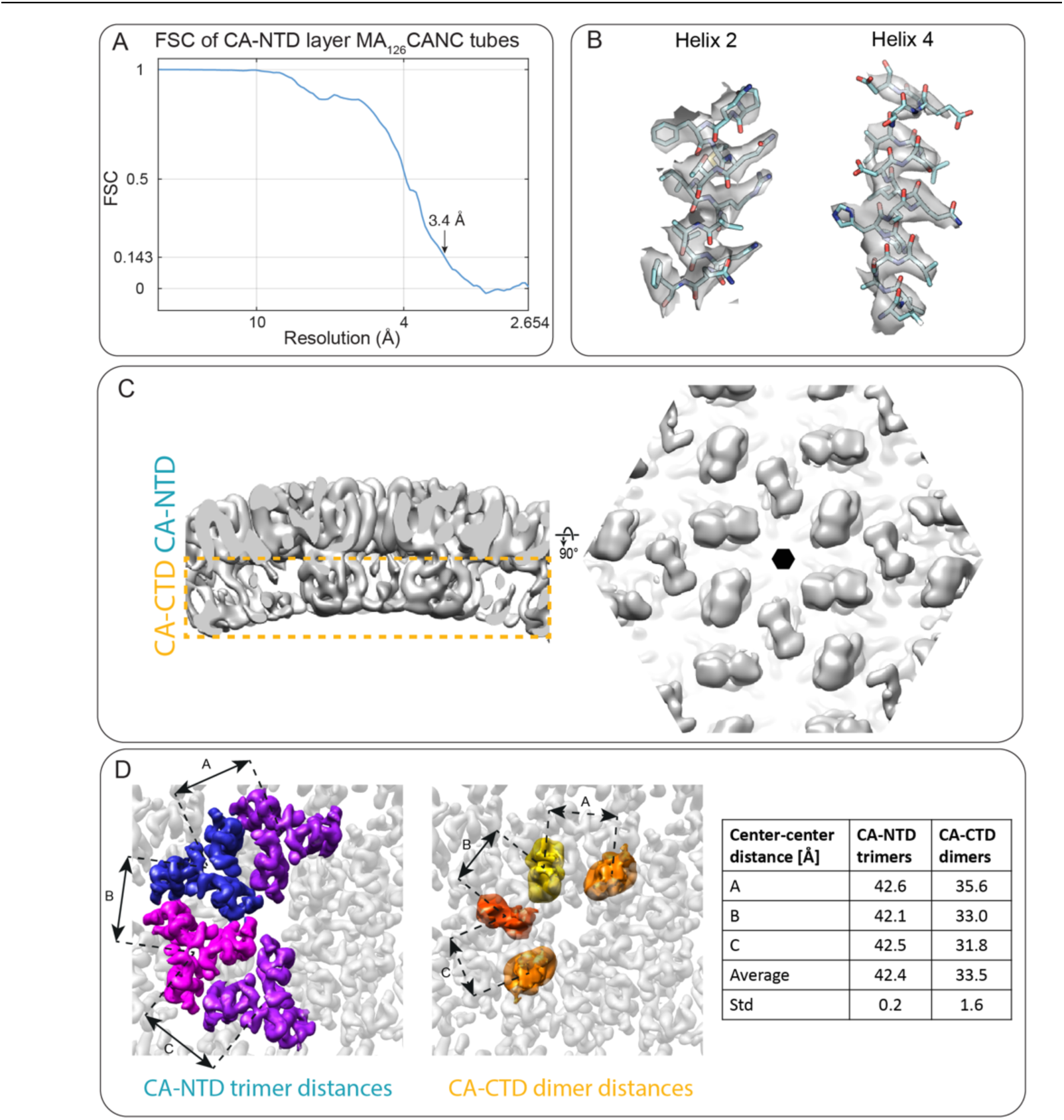
High-resolution cryo-ET structure of HTLV-1 MA126CANC tubes. **A)** FSC for the CA-NTD from MA126CANC tubes, showing a resolution of 3.4 Å at the 0.143 criterion. **B)** Representative EM densities for the CA-NTD, superimposed with the model refined against the 3.4 Å resolution map. At this resolution, the helical pitch and densities for larger side chains are visible. **C)** Low-pass filtered isosurface representation of the subtomogram average of the C2-symmetric CA hexamer from HTLV-1 MA126CANC tubes, as seen in a side view (left) or from within the tube, showing the CA-CTD. This panel relates to the Figure 2C, which shows the same map at the unfiltered 3.4 Å resolution. The orange dashed rectangle indicates the CA-CTD layer, which is not visible at high resolution in Figure 2C. **D)** Distances between adjacent CA-NTD trimers and CA-CTD dimers.

**Figure S8:**
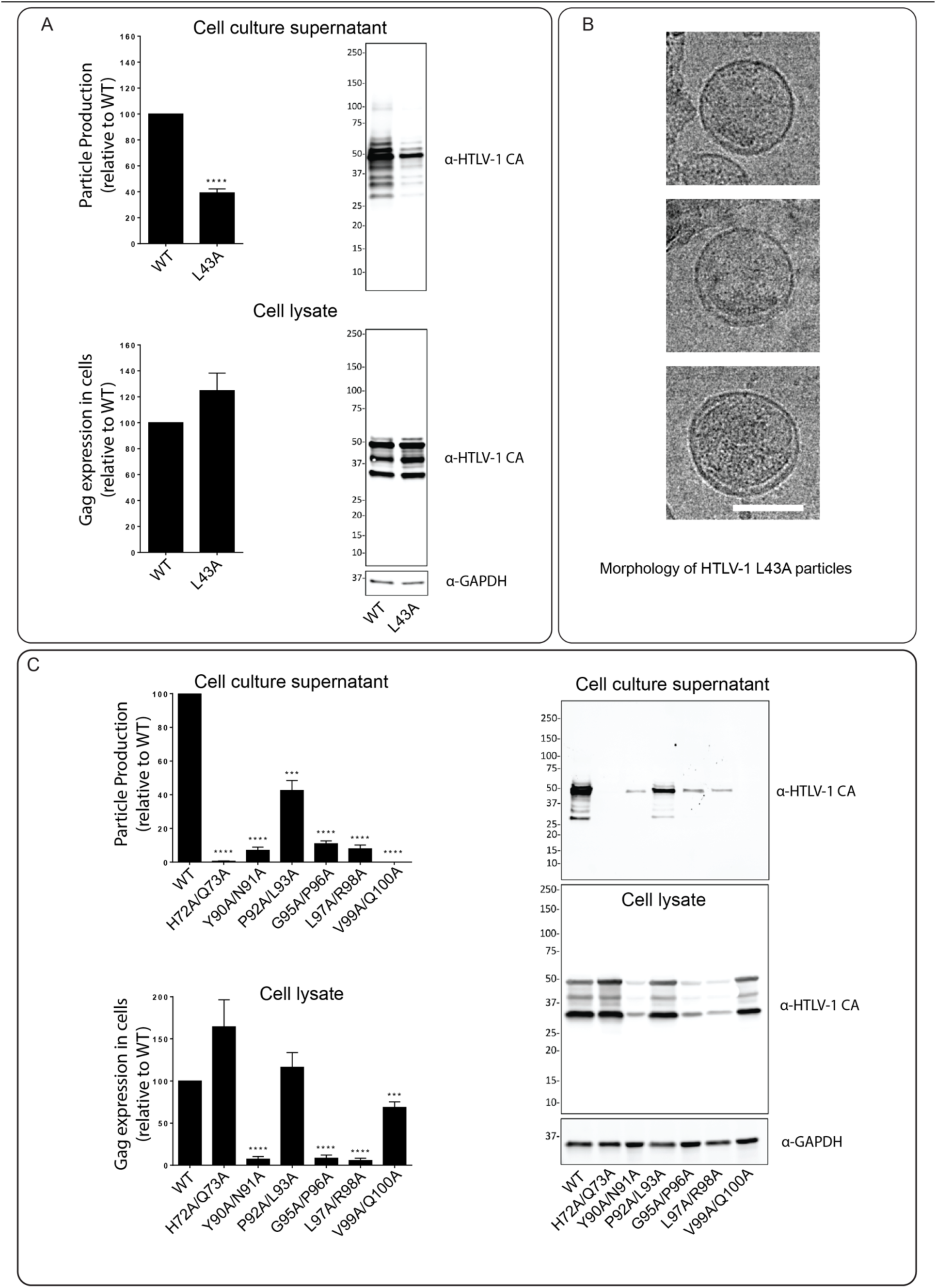
Mutational analysis of the HTLV-1 CA-NTD interfaces. Site-directed mutagenesis of amino acid residues at the CA-NTD interfaces were analyzed for their ability to produce particles relative to that of the WT. Transfection of 293T/17 cells with plasmids expressing the HTLV-1 Gag (WT or alanine-scanning mutant) was done. Cell culture supernatants and cell lysates were harvested 48h post-transfection. **A)** Gag expression in cells and particle production of L43A mutant. Immunoblot analysis was conducted to determine particle production relative to that of WT HTLV-1 Gag. Data was collected from three independent experiments. Error bars represent the standard error of the mean. Significance relative to WT was determined by using an unpaired t-test. ****, *P* < 0.0001. Representative immunoblots of the Gag protein detected from cell culture supernatants and from cell lysates by using an anti-HTLV CA antibody are shown. The locations of the molecular weight markers are indicated along the left side of the immunoblots. **B)** Representative cryo-EM images showing the morphology of the L43A mutant particles. Scale bar = 100 nm. **C)** Gag expression in cells and particle production of the indicated alanine double mutants. Immunoblot analysis was conducted to determine particle production relative to that of WT HTLV-1 Gag. Data was collected from three independent experiments. Error bars represent the standard error of the mean. Significance relative to WT was determined by using an unpaired t-test. ****, *P* < 0.0001; ***, *P* < 0.001; **, *P* < 0.01. Representative immunoblots of the Gag protein detected from cell culture supernatants and from cell lysates by using an anti-HTLV CA antibody are shown.

**Table S1:** Data acquisition statistics.

| Sample |  | HTLV-1 Gag VLPs | HTLV-1<br>MA<sub>126</sub>CANC | HTLV-1<br>MA<sub>126</sub>CANC |
| --- | --- | --- | --- | --- |
| Acquisition settings | Microscope | TFS Titan Krios 3Gi System 1 | TFS Titan Krios 3Gi System 1 | FEI Titan Krios System 2 |
|  | Voltage (keV) | 300 | 300 | 300 |
|  | Detector | Gatan BioQuantum K3 | Gatan BioQuantum K3 | Gatan K2xp |
|  | Energy-filter |  | Yes |  |
|  | Slit width (eV) |  | 20 |  |
|  | Super-resolution mode | Yes | Yes | No |
|  | Å/pixel | 1.381 | 1.381 | 1.327 |
|  | Defocus range (microns) | 1.25-3.5 | 1-3.5 | 1-3 |
|  | Defocus step (microns) |  | 0.25 |  |
|  | Acquisition scheme |  | -60/60°, 3°, Serial EM |  |
|  | Total Dose (electrons/Å<sup>2</sup>) |  | ~145 |  |
|  | Dose rate (eps) | 18 | 21 | 5.4 |
|  | Frame number | 8 | 8 | 10 |
|  | Tomogram number | 85 | 24 | 45 |
| Processing settings | VLPs |  |  |  |
|  | Asymmetric units | 264 000 (CA-NTD)<br>58 000 (CA-CTD) | 268 000 |  |
|  | Final resolution (0.143 FSC) in Å | 5.9 (CA-NTD)<br>6.2 (CA-CTD) | 3.4 |  |

**Table S2:**
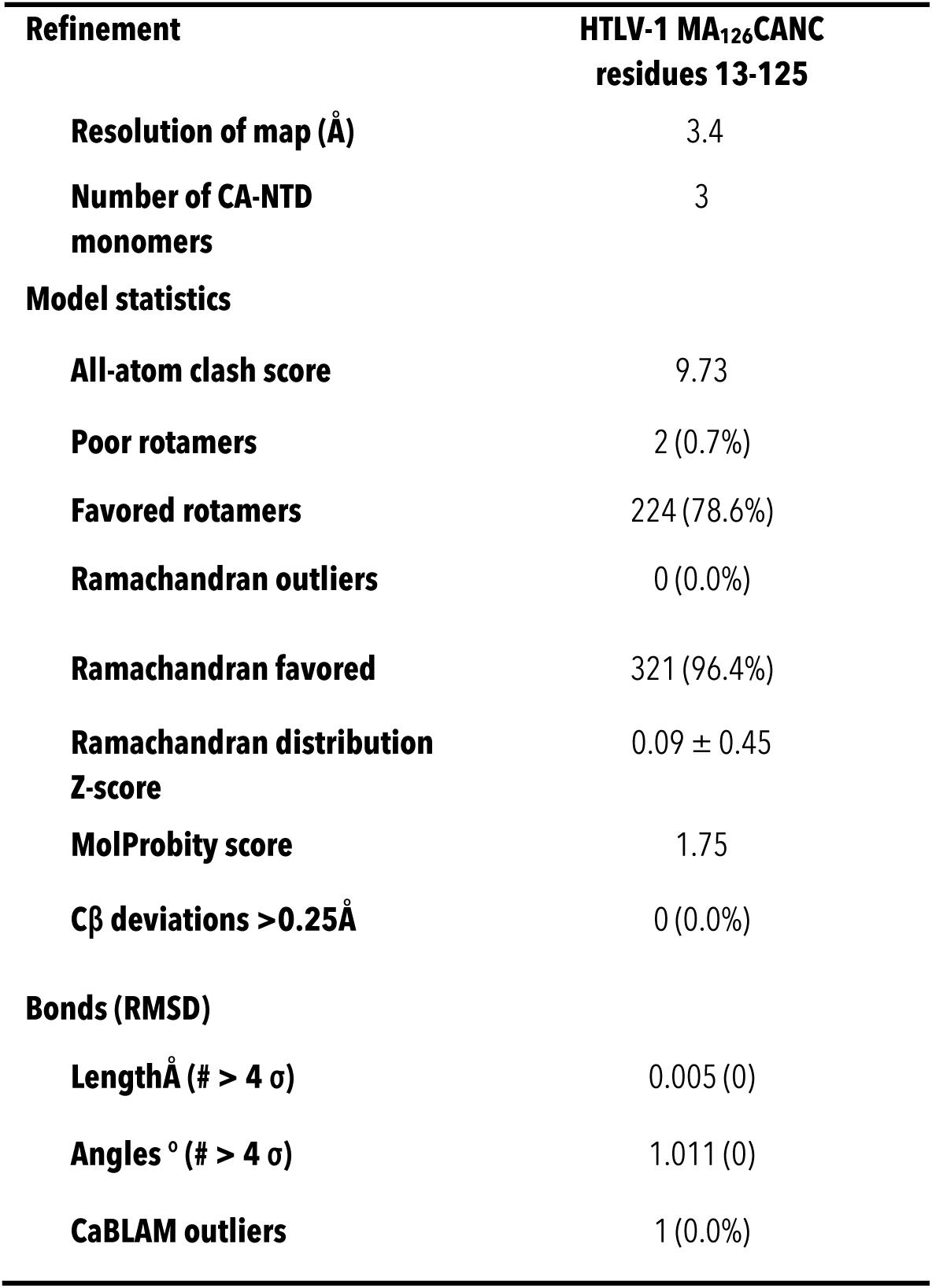
Model refinement statistics.

| Refinement | HTLV-1 MA<sub>126</sub>CANC<br>residues 13-125 |
| --- | --- |
| Resolution of map (Å) | 3.4 |
| Number of CA-NTD<br>monomers | 3 |
| **Model statistics** |  |
| All-atom clash score | 9.73 |
| Poor rotamers | 2 (0.7%) |
| Favored rotamers | 224 (78.6%) |
| Ramachandran outliers | 0 (0.0%) |
| Ramachandran favored | 321 (96.4%) |
| Ramachandran distribution<br>Z-score | 0.09 ± 0.45 |
| MolProbity score | 1.75 |
| Cβ deviations >0.25Å | 0 (0.0%) |
| **Bonds (RMSD)** |  |
| LengthÅ (# > 4 $\sigma$) | 0.005 (0) |
| Angles ° (# > 4 $\sigma$) | 1.011 (0) |
| CaBLAM outliers | 1 (0.0%) |

## Supplementary Movie Legends

**Movie S1**: Cryo-electron tomogram of HTLV-1 Gag-based VLPs.

**Movie S2:** Tour through the immature HTLV-1 CA lattice, annotating the CA-NTD and CA-CTD interfaces as described in Figure 1.

**Movie S3:** Cryo-electron tomogram of HTLV-1 MA126CANC tubes.

## References

1. A. Gessain, A. Gessain, O. Cassar, Epidemiological Aspects and World Distribution of HTLV-1 Infection. Frontiers in Microbiology. 3 (2012) (available at https://www.frontiersin.org/articles/10.3389/fmicb.2012.00388).

2. L. B. Cook, M. Elemans, A. G. Rowan, B. Asquith, HTLV-1: Persistence and pathogenesis. Virology. 435, 131–140 (2013).

3. R. Mahieux, A. Gessain, Adult T-cell leukemia/lymphoma and HTLV-1. Current Hematologic Malignancy Reports. 2, 257–264 (2007).

4. E. O. Freed, HIV-1 assembly, release and maturation. Nature Reviews Microbiology. 13, 484–496 (2015).

5. D. Ako-Adjei, M. C. Johnson, V. M. Vogt, The Retroviral Capsid Domain Dictates Virion Size, Morphology, and Coassembly of Gag into Virus-Like Particles. Journal of Virology. 79, 13463–13472 (2005).

6. S. Mattei, F. K. M. Schur, J. A. Briggs, Retrovirus maturation - An extraordinary structural transformation. Current Opinion in Virology. 18, 27–35 (2016).

7. D. L. Bush, V. M. Vogt, In Vitro Assembly of Retroviruses. Annual Review of Virology. 1, 561–580 (2014).

8. F. K. M. Schur, W. J. H. Hagen, M. Rumlová, T. Ruml, B. Müller, H. G. Kraüsslich, J. A. G. Briggs, Structure of the immature HIV-1 capsid in intact virus particles at 8.8 Å resolution. Nature. 517, 505–508 (2015).

9. F. K. M. Schur, M. Obr, W. J. H. Hagen, W. Wan, A. J. Jakobi, J. M. Kirkpatrick, C. Sachse, H. G. Kräusslich, J. A. G. Briggs, An atomic model of HIV-1 capsid-SP1 reveals structures regulating assembly and maturation. Science. 353, 506–508 (2016).

10. G. Zhao, J. R. Perilla, E. L. Yufenyuy, X. Meng, B. Chen, J. Ning, J. Ahn, A. M. Gronenborn, K. Schulten, C. Aiken, P. Zhang, Mature HIV-1 capsid structure by cryo-electron microscopy and all-atom molecular dynamics. Nature. 497, 643–646 (2013).

11. R. T. Schirra, N. F. B. dos Santos, K. K. Zadrozny, I. Kucharska, B. K. Ganser-Pornillos, O. Pornillos, A molecular switch modulates assembly and host factor binding of the HIV-1 capsid. Nat Struct Mol Biol, 1–8 (2023).

12. S. Khorasanizadeh, R. Campos-Olivas, C. Clark, M. Summers, Sequence-specific 1H, 13C and 15N chemical shift assignment and secondary structure of the HTLV-I capsid protein. Journal of Biomolecular NMR. 14, 199–200 (1999).

13. C. Tang, Y. Ndassa, M. F. Summers, Structure of the N-terminal 283-residue fragment of the immature HIV-1 Gag polyprotein. Nat Struct Biol. 9, 537–543 (2002).

14. V. Bartonova, S. Igonet, J. Sticht, B. Glass, A. Habermann, M. C. Vaney, P. Sehr, J. Lewis, F. A. Rey, H. G. Krausslich, Residues in the HIV-1 capsid assembly inhibitor binding site are essential for maintaining the assembly-competent quaternary structure of the capsid protein. J Biol Chem. 283, 32024–32033 (2008).

15. R. L. Kingston, T. Fitzon-Ostendorp, E. Z. Eisenmesser, G. W. Schatz, V. M. Vogt, C. B. Post, M. G. Rossmann, Structure and self-association of the Rous sarcoma virus capsid protein. Structure. 8, 617–628 (2000).

16. R. Campos-Olivas, J. L. Newman, M. F. Summers, Solution structure and dynamics of the Rous sarcoma virus capsid protein and comparison with capsid proteins of other retroviruses. J Mol Biol. 296, 633–649 (2000).

17. C. C. Cornilescu, F. Bouamr, X. Yao, C. Carter, N. Tjandra, Structural analysis of the N-terminal domain of the human T-cell leukemia virus capsid protein. J Mol Biol. 306, 783–797 (2001).

18. K. Qu, B. Glass, M. Doležal, F. K. M. Schur, B. Murciano, A. Rein, M. Rumlová, T. Ruml, H.-G. Kräusslich, J. A. G. Briggs, Structure and architecture of immature and mature murine leukemia virus capsids. Proceedings of the National Academy of Sciences. 115, E11751– E11760 (2018).

19. F. Mammano, A. Ohagen, S. Höglund, H. G. Göttlinger, Role of the major homology region of human immunodeficiency virus type 1 in virion morphogenesis. Journal of Virology. 68, 4927–4936 (1994).

20. T. A. M. Bharat, N. E. Davey, P. Ulbrich, J. D. Riches, A. de Marco, M. Rumlova, C. Sachse, T. Ruml, J. A. G. Briggs, Structure of the immature retroviral capsid at 8 Å resolution by cryo-electron microscopy. Nature. 487, 385–389 (2012).

21. K. Strohalmová-Bohmová, V. Spiwok, M. Lepšík, R. Hadravová, I. Křížová, P. Ulbrich, I. Pichová, L. Bednárová, T. Ruml, M. Rumlová, Role of Mason-Pfizer Monkey Virus CA-NC Spacer Peptide-Like Domain in Assembly of Immature Particles. J Virol. 88, 14148–14160 (2014).

22. O. Pornillos, B. K. Ganser-Pornillos, Maturation of retroviruses. Current Opinion in Virology. 36, 47–55 (2019).

23. M. A. Accola, B. Strack, H. G. Göttlinger, Efficient Particle Production by Minimal Gag Constructs Which Retain the Carboxy-Terminal Domain of Human Immunodeficiency Virus Type 1 Capsid-p2 and a Late Assembly Domain. Journal of Virology. 74, 5395–5402 (2000).

24. A. Borsetti, Å. Öhagen, H. G. Göttlinger, The C-Terminal Half of the Human Immunodeficiency Virus Type 1 Gag Precursor Is Sufficient for Efficient Particle Assembly. Journal of Virology. 72, 9313–9317 (1998).

25. F. K. M. Schur, R. A. Dick, W. J. H. Hagen, V. M. Vogt, J. A. G. Briggs, The Structure of Immature Virus-Like Rous Sarcoma Virus Gag Particles Reveals a Structural Role for the p10 Domain in Assembly. Journal of Virology. 89, 10294–10302 (2015).

26. R. A. Dick, C. Xu, D. R. Morado, V. Kravchuk, C. L. Ricana, T. D. Lyddon, A. M. Broad, J. R. Feathers, M. C. Johnson, V. M. Vogt, J. R. Perilla, J. A. G. Briggs, F. K. M. Schur, Structures of immature EIAV Gag lattices reveal a conserved role for IP6 in lentivirus assembly. PLoS Pathogens. 16 (2020), doi:10.1371/journal.ppat.1008277.

27. J. Temple, T. N. Tripler, Q. Shen, Y. Xiong, A snapshot of HIV-1 capsid–host interactions. Current Research in Structural Biology. 2, 222– 228 (2020).

28. F. Rayne, F. Bouamr, J. Lalanne, R. Z. Mamoun, The N-Terminal Domain of the Human T-Cell Leukemia Virus Type 1 Capsid Protein Is Involved in Particle Formation. Journal of Virology. 75, 5277–5287 (2001).

29. S. Cao, J. O. Maldonado, I. F. Grigsby, L. M. Mansky, W. Zhang, Analysis of human T-cell leukemia virus type 1 particles by using cryo-electron tomography. Journal of virology. 89, 2430–5 (2015).

30. J. O. Maldonado, S. Cao, W. Zhang, L. M. Mansky, Distinct morphology of human T-cell leukemia virus type 1-like particles. Viruses. 8, 132 (2016).

31. J. L. Martin, S. Cao, J. O. Maldonado, W. Zhang, L. M. Mansky, Distinct Particle Morphologies Revealed through Comparative Parallel Analyses of Retrovirus-Like Particles. Journal of Virology. 90, 8074–8084 (2016).

32. S. K. Carnes, J. H. Sheehan, C. Aiken, Inhibitors of the HIV-1 capsid, a target of opportunity. Current Opinion in HIV and AIDS. 13, 359 (2018).

33. E. R. Wright, J. B. Schooler, H. J. Ding, C. Kieffer, C. Fillmore, W. I. Sundquist, G. J. Jensen, Electron cryotomography of immature HIV-1 virions reveals the structure of the CA and SP1 Gag shells. EMBO Journal. 26, 2218–2226 (2007).

34. D. Tegunov, L. Xue, C. Dienemann, P. Cramer, J. Mahamid, Multi-particle cryo-EM refinement with M visualizes ribosome-antibiotic complex at 3.5 Å in cells. Nature Methods. 18, 186–193 (2021).

35. J. M. Wagner, K. K. Zadrozny, J. Chrustowicz, M. D. Purdy, M. Yeager, B. K. Ganser-Pornillos, O. Pornillos, Crystal structure of an HIV assembly and maturation switch. eLife. 5 (2016), doi:10.7554/eLife.17063.

36. T. A. M. Bharat, L. R. Castillo Menendez, W. J. H. Hagen, V. Lux, S. Igonet, M. Schorb, F. K. M. Schur, H.-G. Krausslich, J. A. G. Briggs, Cryo-electron microscopy of tubular arrays of HIV-1 Gag resolves structures essential for immature virus assembly. Proceedings of the National Academy of Sciences. 111, 8233–8238 (2014).

37. J. L. Martin, L. M. Mendonça, R. Marusinec, J. Zuczek, I. Angert, R. J. Blower, J. D. Mueller, J. R. Perilla, W. Zhang, L. M. Mansky, Critical Role of the Human T-Cell Leukemia Virus Type 1 Capsid N-Terminal Domain for Gag-Gag Interactions and Virus Particle Assembly. Journal of Virology. 92, e00333–18 (2018).

38. A. Real-Hohn, M. Groznica, N. Löffler, D. Blaas, H. Kowalski, nanoDSF: In vitro Label-Free Method to Monitor Picornavirus Uncoating and Test Compounds Affecting Particle Stability. Frontiers in Microbiology. 11 (2020) (available at https://www.frontiersin.org/articles/10.3389/fmicb.2020.01442).

39. J. A. G. Briggs, M. C. Johnson, M. N. Simon, S. D. Fuller, V. M. Vogt, Cryo-electron Microscopy Reveals Conserved and Divergent Features of Gag Packing in Immature Particles of Rous Sarcoma Virus and Human Immunodeficiency Virus. J Mol Biol. 355, 157–168 (2006).

40. J. L. Martin, L. M. Mendonça, I. Angert, J. D. Mueller, W. Zhang, L. M. Mansky, Disparate Contributions of Human Retrovirus Capsid Subdomains to Gag-Gag Oligomerization, Virus Morphology, and Particle Biogenesis. Journal of Virology. 91, e00298–17 (2017).

41. M. A. Accola, S. Höglund, H. G. Göttlinger, A Putative **α**-Helical Structure Which Overlaps the Capsid-p2 Boundary in the Human Immunodeficiency Virus Type 1 Gag Precursor Is Crucial for Viral Particle Assembly. Journal of Virology. 72, 2072–2078 (1998).

42. S. R. Cheslock, D. T. K. Poon, W. Fu, T. D. Rhodes, L. E. Henderson, K. Nagashima, C. F. McGrath, W.-S. Hu, Charged Assembly Helix Motif in Murine Leukemia Virus Capsid: an Important Region for Virus Assembly and Particle Size Determination. Journal of Virology. 77, 7058–7066 (2003).

43. U. K. von Schwedler, K. M. Stray, J. E. Garrus, W. I. Sundquist, Functional Surfaces of the Human Immunodeficiency Virus Type 1 Capsid Protein. Journal of Virology. 77, 5439–5450 (2003).

44. T. Füzik, R. Píchalová, F. K. M. Schur, K. Strohalmová, I. Křížová, R. Hadravová, M. Rumlová, J. A. G. Briggs, P. Ulbrich, T. Ruml, Nucleic Acid Binding by Mason-Pfizer Monkey Virus CA Promotes Virus Assembly and Genome Packaging. Journal of Virology. 90, 4593–4603 (2016).

45. D. F. Qualley, K. M. Stewart-Maynard, F. Wang, M. Mitra, R. J. Gorelick, I. Rouzina, M. C. Williams, K. Musier-Forsyth, C-terminal Domain Modulates the Nucleic Acid Chaperone Activity of Human T-cell Leukemia Virus Type 1 Nucleocapsid Protein via an Electrostatic Mechanism*. Journal of Biological Chemistry. 285, 295–307 (2010).

46. J. E. Eschbach, J. L. Elliott, W. Li, K. K. Zadrozny, K. Davis, S. J. Mohammed, D. Q. Lawson, O. Pornillos, A. N. Engelman, S. B. Kutluay, Capsid Lattice Destabilization Leads to Premature Loss of the Viral Genome and Integrase Enzyme during HIV-1 Infection. Journal of Virology. 95, e00984–20 (2020).

47. R. A. Dick, K. K. Zadrozny, C. Xu, F. K. M. Schur, T. D. Lyddon, C. L. Ricana, J. M. Wagner, J. R. Perilla, B. K. Ganser-Pornillos, M. C. Johnson, O. Pornillos, V. M. Vogt, Inositol phosphates are assembly co-factors for HIV-1. Nature. 560, 509–512 (2018).

48. S. M. Bester, G. Wei, H. Zhao, D. Adu-Ampratwum, N. Iqbal, V. V. Courouble, A. C. Francis, A. S. Annamalai, P. K. Singh, N. Shkriabai, P. Van Blerkom, J. Morrison, E. M. Poeschla, A. N. Engelman, G. B. Melikyan, P. R. Griffin, J. R. Fuchs, F. J. Asturias, M. Kvaratskhelia, Structural and mechanistic bases for a potent HIV-1 capsid inhibitor. Science. 370, 360–364 (2020).

49. J. O. Link, M. S. Rhee, W. C. Tse, J. Zheng, J. R. Somoza, W. Rowe, R. Begley, A. Chiu, A. Mulato, D. Hansen, E. Singer, L. K. Tsai, R. A. Bam, C.-H. Chou, E. Canales, G. Brizgys, J. R. Zhang, J. Li, M. Graupe, P. Morganelli, Q. Liu, Q. Wu, R. L. Halcomb, R. D. Saito, S. D. Schroeder, S. E. Lazerwith, S. Bondy, D. Jin, M. Hung, N. Novikov, X. Liu, A. G. Villaseñor, C. E. Cannizzaro, E. Y. Hu, R. L. Anderson, T. C. Appleby, B. Lu, J. Mwangi, A. Liclican, A. Niedziela-Majka, G. A. Papalia, M. H. Wong, S. A. Leavitt, Y. Xu, D. Koditek, G. J. Stepan, H. Yu, N. Pagratis, S. Clancy, S. Ahmadyar, T. Z. Cai, S. Sellers, S. A. Wolckenhauer, J. Ling, C. Callebaut, N. Margot, R. R. Ram, Y.-P. Liu, R. Hyland, G. I. Sinclair, P. J. Ruane, G. E. Crofoot, C. K. McDonald, D. M. Brainard, L. Lad, S. Swaminathan, W. I. Sundquist, R. Sakowicz, A. E. Chester, W. E. Lee, E. S. Daar, S. R. Yant, T. Cihlar, Clinical targeting of HIV capsid protein with a long-acting small molecule. Nature. 584, 614–618 (2020).

50. C. S. Adamson, K. Salzwedel, E. O. Freed, Virus maturation as a new HIV-1 therapeutic target. Expert Opinion on Therapeutic Targets. 13, 895–908 (2009).

51. D. N. Mastronarde, “SerialEM: A program for automated tilt series acquisition on Tecnai microscopes using prediction of specimen position” in Microscopy and Microanalysis (Cambridge University Press, 2003), vol. 9, pp. 1182–1183.

52. W. J. H. Hagen, W. Wan, J. A. G. Briggs, Implementation of a cryo-electron tomography tilt-scheme optimized for high resolution subtomogram averaging. Journal of Structural Biology. 197, 191–198 (2017).

53. T. Grant, N. Grigorieff, Measuring the optimal exposure for single particle cryo-EM using a 2.6 Å reconstruction of rotavirus VP6. eLife. 4, e06980 (2015).

54. J. R. Kremer, D. N. Mastronarde, J. R. McIntosh, Computer Visualization of Three-Dimensional Image Data Using IMOD. J Struct Biol. 116, 71–76 (1996).

55. M. Obr, C. L. Ricana, N. Nikulin, J.-P. R. Feathers, M. Klanschnig, A. Thader, M. C. Johnson, V. M. Vogt, F. K. M. Schur, R. A. Dick, Structure of the mature Rous sarcoma virus lattice reveals a role for IP6 in the formation of the capsid hexamer. Nature Communications. 12, 3226 (2021).

56. B. Turoňová, F. K. M. Schur, W. Wan, J. A. G. Briggs, Efficient 3D-CTF correction for cryo-electron tomography using NovaCTF improves subtomogram averaging resolution to 3.4 Å. Journal of Structural Biology. 199, 187–195 (2017).

57. D. Castano-Diez, M. Kudryashev, M. Arheit, H. Stahlberg, Dynamo: a flexible, user-friendly development tool for subtomogram averaging of cryo-EM data in high-performance computing environments. J Struct Biol. 178, 139–151 (2012).

58. D. Tegunov, P. Cramer, Real-time cryo-electron microscopy data preprocessing with Warp. Nature Methods. 16, 1146–1152 (2019).

59. S. H. W. Scheres, RELION: Implementation of a Bayesian approach to cryo-EM structure determination. J Struct Biol. 180, 519–530 (2012).

60. M. Mirdita, K. Schütze, Y. Moriwaki, L. Heo, S. Ovchinnikov, M. Steinegger, ColabFold: making protein folding accessible to all. Nat Methods. 19, 679–682 (2022).

61. E. F. Pettersen, T. D. Goddard, C. C. Huang, G. S. Couch, D. M. Greenblatt, E. C. Meng, T. E. Ferrin, UCSF Chimera--a visualization system for exploratory research and analysis. J Comput Chem. 25, 1605–1612 (2004).

62. M. Obr, W. J. H. Hagen, R. A. Dick, L. Yu, A. Kotecha, F. K. M. Schur, Exploring high-resolution cryo-ET and subtomogram averaging capabilities of contemporary DEDs. Journal of Structural Biology. 214, 107852 (2022).

63. T. I. Croll, ISOLDE: a physically realistic environment for model building into low-resolution electron-density maps. Acta Crystallographica Section D. 74, 519–530 (2018).

64. T. D. Goddard, C. C. Huang, E. C. Meng, E. F. Pettersen, G. S. Couch, J. H. Morris, T. E. Ferrin, UCSF ChimeraX: Meeting modern challenges in visualization and analysis. Protein Science. 27, 14–25 (2018).

65. P. D. Adams, P. V. Afonine, G. Bunkoczi, V. B. Chen, I. W. Davis, N. Echols, J. J. Headd, L.-W. Hung, G. J. Kapral, R. W. Grosse-Kunstleve, A. J. McCoy, N. W. Moriarty, R. Oeffner, R. J. Read, D. C. Richardson, J. S. Richardson, T. C. Terwilliger, P. H. Zwart, PHENIX: a comprehensive Python-based system for macromolecular structure solution. Acta Crystallographica Section D. 66, 213–221 (2010).

66. P. Emsley, K. Cowtan, Coot: model-building tools for molecular graphics. Acta Crystallographica Section D. 60, 2126–2132 (2004).

67. Schrodinger LLC, “The PyMOL Molecular Graphics System, Version 1.3r1” (2010).

68. F. M. Heydenreich, T. Miljuš, R. Jaussi, R. Benoit, D. Milić, D. B. Veprintsev, High-throughput mutagenesis using a two-fragment PCR approach. Sci Rep. 7, 6787 (2017).

